# Discovery of a Novel, Selective and Short-Acting Skeletal Myosin II Inhibitor

**DOI:** 10.1101/2020.07.28.225383

**Authors:** Laszlo Radnai, Matthew Surman, Madalyn Hafenbreidel, Erica J. Young, Rebecca F. Stremel, Li Lin, Paolo Pasetto, Xiaomin Jin, Mackenzie Geedy, Joni-Rae Partridge, Aagam Patel, Michael Conlon, James R. Sellers, Michael D. Cameron, Gavin Rumbaugh, Patrick R. Griffin, Theodore M. Kamenecka, Courtney A. Miller

## Abstract

Myosin IIs, actin-based motors that utilize the chemical energy of ATP to generate force, have potential as therapeutic targets. Their heavy chains differentiate the family into muscle (skeletal [SkMII], cardiac, smooth) and nonmuscle myosin IIs. Despite therapeutic potential for muscle disorders, no SkMII-specific inhibitor has been reported and characterized. Here we present the discovery, synthesis and characterization of “skeletostatins”, novel derivatives of the pan-myosin II inhibitor blebbistatin, with selectivity within the myosin IIs for SkMII. In addition, the skeletostatins bear improved potency, solubility and photostability, without cytotoxicity. Based on its optimal *in vitro* profile, Skeletostatin 1’s *in vivo* tolerability, efficacy and pharmacokinetics were determined. Skeletostatin 1 was well-tolerated in mice, impaired motor performance, and had an excellent muscle to plasma ratio. Skeletostatins are useful probes for basic research and a strong starting point for drug development.

## Introduction

Myosins are intracellular actin-based motor proteins that convert the chemical energy stored in ATP to mechanical work^[1]^. A recent phylogenetic analysis of more than 4,000 myosin gene sequences from 340 diverse eukaryote species identified 45 myosin classes within the myosin superfamily^[2]^. Different myosins have been adapted to various roles during evolution, including muscle contraction, cell locomotion, cell division, and intracellular trafficking or anchoring of organelles and cargoes^[1]^. In humans, 38 myosin genes belonging to 12 classes are expressed in a cell type-specific manner. Myosin IIs, also known as conventional myosins, consist of two heavy chains, which bear the force generating motor heads, and several light chains^[1a]^. Myosin II family members are distinguished by their different heavy chains, which results in skeletal (SkMII), cardiac (CMII) and smooth (SmMII) muscle myosin IIs, as well as nonmuscle myosin IIs (NMII)^[1a]^.

Blebbistatin (blebb) was discovered in a high-throughput screening assay targeting NMII and used to conclusively demonstrate myosin II’s integral role in cytokinesis^[3]^. Blebb inhibits the steady state ATPase activity of myosin by binding to the myosin-ADP-inorganic phosphate complex and blocking inorganic phosphate release, thereby trapping myosin in a weak actin binding state^[4]^. The activity and specificity of blebb has been relatively well-characterized. It inhibits both the ATPase and force generation activities of SkMII, CMII, NMII and, to a lesser degree, SmMII, but not other myosin superfamily members from classes I, V, and X^[3, 5]^. Accordingly, it blocks not only NMII-related functions^[3]^, but also skeletal^[6]^, cardiac^[6a, 7]^ and smooth^[8]^ muscle functions. Blebb is most potent against SkMII and has long been recognized as a potential starting point for studies aiming to develop novel drug candidates for the treatment of various conditions^[9]^.

Limited SAR has been performed on blebb, exploring modifications to the A, B, C and D rings (**Scheme 1**). Modifications to the A-ring resulted in compounds with reduced potency against SkMII^[10]^ and C-ring modifications rendered the compounds inactive at SkMII^[11]^. D-ring substitutions improved blebb’s physicochemical properties, particularly photostability, solubility, and fluorescence ^[12]^. However, to date, no attempt has been made to alter blebb’s selectivity profile within the myosin II family. Here we set out to determine if blebb could be modified to improve its selectivity for SkMII over the other myosin II’s. Given the critical role of CMII to cardiomyocyte contractility^[13]^ and, therefore, heart function, we hypothesized that derivatives that retained SkMII inhibition, but reduced CMII inhibition, would result in safer probes for assessing *in vivo* functions, but also as optimal starting points in future studies aimed at developing drug candidates for the treatment of various SkMII-related conditions.

## Results and Discussion

### Synthesis, purification and analysis of skeletostatins

The synthesis of (*S*)-3a-hydroxy-6-methyl-1-(2-methyloxazolo[4,5-*b*]pyridin-6-yl)-1,2,3,3a-tetrahydro-4*H*-pyrrolo[2,3-*b*]quinolin-4-one, (*S*)-N-(4-(3a-hydroxy-6-methyl-4-oxo-2,3,3a,4-tetrahydro-1*H*-pyrrolo[2,3-*b*]quinolin-1-yl)phenyl)acetamide and (*S*)-3a-hydroxy-6-methyl-1-(4-morpholinophenyl)-1,2,3,3a-tetrahydro-4*H*-pyrrolo[2,3-*b*]quinolin-4-one was performed. These compounds were named Skeletostatin 1, 2 and 3, respectively (**Scheme 1**). In order to compare these newly synthesized compounds to the parent compound, (*S*)-blebb was also synthesized. (See Supporting Information for the details of chemical synthesis.)

**Scheme 1.**
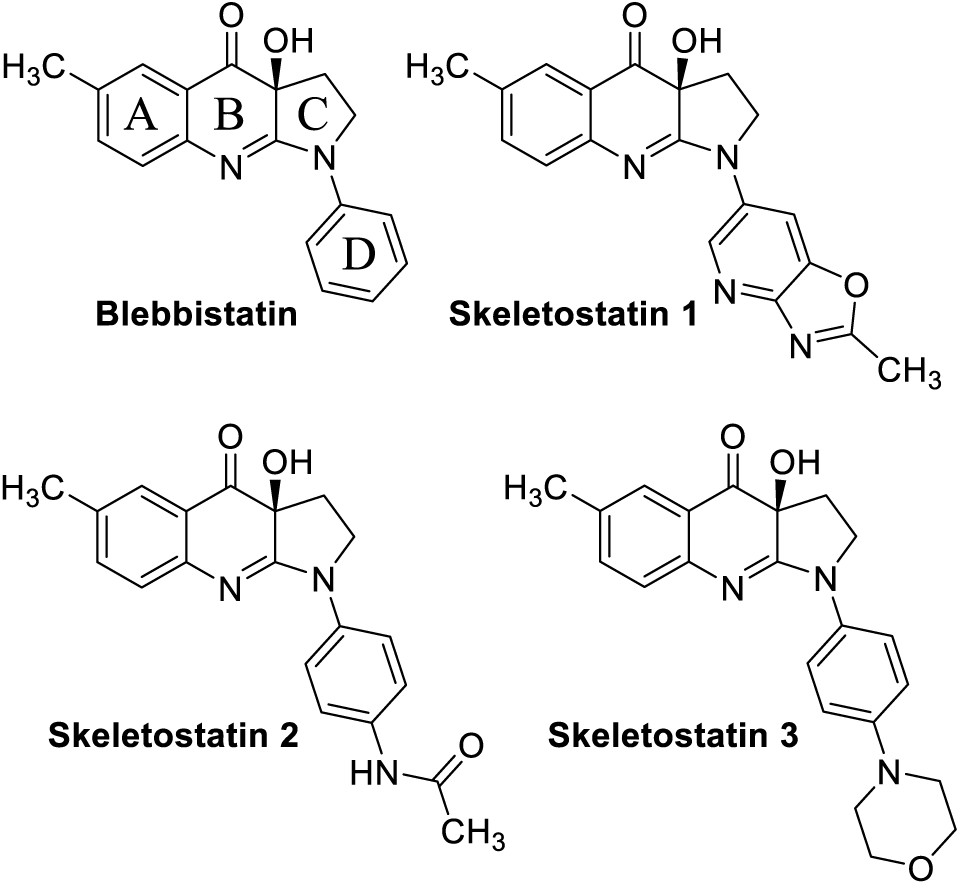
Chemical structures of blebbistatin and skeletostatins.

Skeletostatin 1 was prepared using a modification of the procedure reported by Lawson, et al. (**Scheme 2**)^[14]^. Pyrrolidinone **3** was obtained via a CuI-catalyzed *N*-arylation of 2-pyrrolidinone (**2**) with 1-iodo-4-methoxybenzene (**1**). Treatment of pyrrolidinone **3** with POCl_3_ and aniline **4** provided amidine **5**. Cyclization was accomplished with an excess of LiHMDS to give quinolinone **6**. Asymmetric hydroxylation with oxaziridine **7** provided (*S*)-4’-methoxyblebbistatin (**8**)^[15]^. TIPS protection of the tertiary alcohol followed by *p*-methoxyphenyl removal with CAN provided NH amidine **10**. Installation of the methyloxazolopyridine ring was accomplished with a palladium-catalyzed coupling using Pd_2_(dba)_3_ and Xantphos ligand. A final TIPS deprotection provided Skeletostatin 1.

**Scheme 2.**
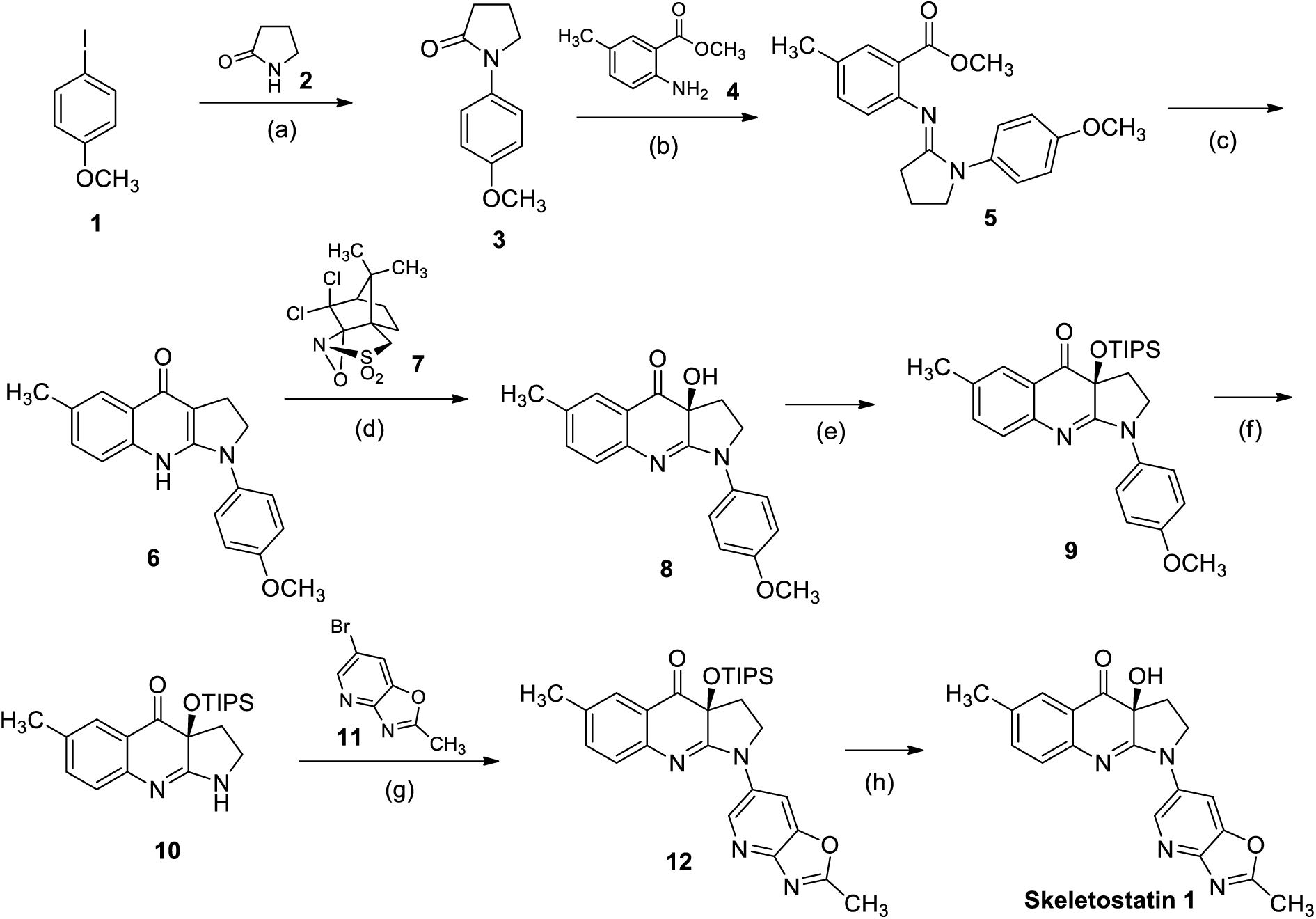
Synthesis of **Skeletostatin 1**. Reagents and conditions: (a) **2**, CuI, Cs_2_CO_3_, and *N,N*’-dimethyl-(1*R*,2*R*)-1,2-cyclohexanediamine, toluene, reflux, 16 h, 86%; (b) **4**, POCl_3_, CH_2_Cl_2_, reflux, 16 h, 60%; (c) LiHMDS, THF, 0 °C, 2 h, 59%; (d) **7**, LiHMDS, THF –20 to 0 °C, 1 h, 96%; (e) TIPSOTf, DIPEA, CH_2_Cl_2_, reflux, 16 h, 43%; (f) CAN, CH_3_CN, H_2_O, 0 °C, 2 h, 99%; (g) Pd_2_(dba)_3_, XANTPHOS, Cs_2_CO_3_, 1,4-dioxane, 120 °C, 16 h, 71%; (h) TBAF, THF, rt, 1 h, 24%

The preparation of Skeletostatin 2 commenced with the CuI-catalyzed *N*-arylation of 2-pyrrolidinone (**2**) with 1-bromo-4-iodobenzene (**13**) to give pyrrolidinone **14** (**Scheme 3**). Amidine **15** was prepared from pyrrolidinone **14** and aniline **4** in the presence of POCl_3_. Cyclization of **15** and asymmetric hydroxylation were carried out in a single pot^[12d]^ to provide 4’-bromoblebbistatin **16**^[10]^. A second CuI-catalyzed *N*-arylation was used to install the acetamide group to give Skeletostatin 2.

**Scheme 3.**
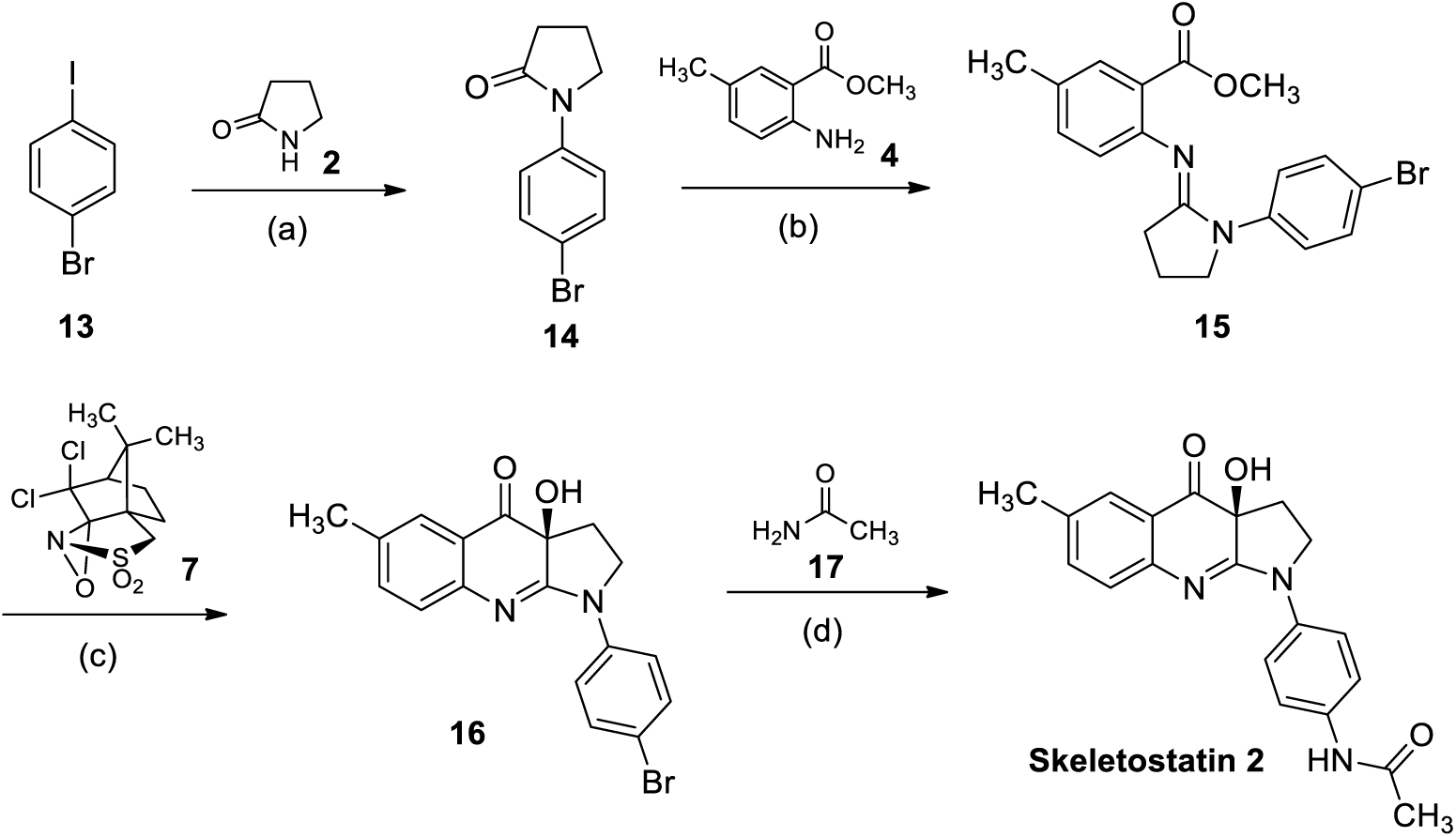
Synthesis of **Skeletostatin 2**. Reagents and conditions: (a) **2**, CuI, Cs_2_CO_3_, and *N,N*’-dimethyl-(1*R*,2*R*)-1,2-cyclohexanediamine, DMSO, 110 °C, 16 h, 75%; (b) **4**, POCl_3_, CH_2_Cl_2_, 45 °C, 96 h, 38%; (c) **7**, LiHMDS, THF, –78 to 0 °C, 2 h, 36%; (e) CuI, K_2_CO_3_, and *N,N*’-dimethyl-(1*R*,2*R*)-1,2-cyclohexanediamine, 1,4-dioxane, 100 °C, 16 h, 28%

A modified route was developed for the synthesis of Skeletostatin 3 (**Scheme 4**). Unlike the synthesis of Skeletostatin 1, which employed a late stage intermediate for elaboration (amidine **10**) that contained the chiral center, the late stage intermediate (amidine **21**) in the synthesis of Skeletostatin 3 was achiral. The chiral center was established in the final step, thus conserving the expensive oxaziridine reagent. After formation of amidine **19** and cyclization to give quinolinol **20**, the benzyl group was removed with AlCl_3_ to give amidine **21**. Compound **21** was elaborated with 4-(4-bromophenyl)morpholine (**22**) through a copper catalyzed coupling to give quinolinol **23**. Asymmetric hydroxylation provided Skeletostatin 3.

**Scheme 4.**
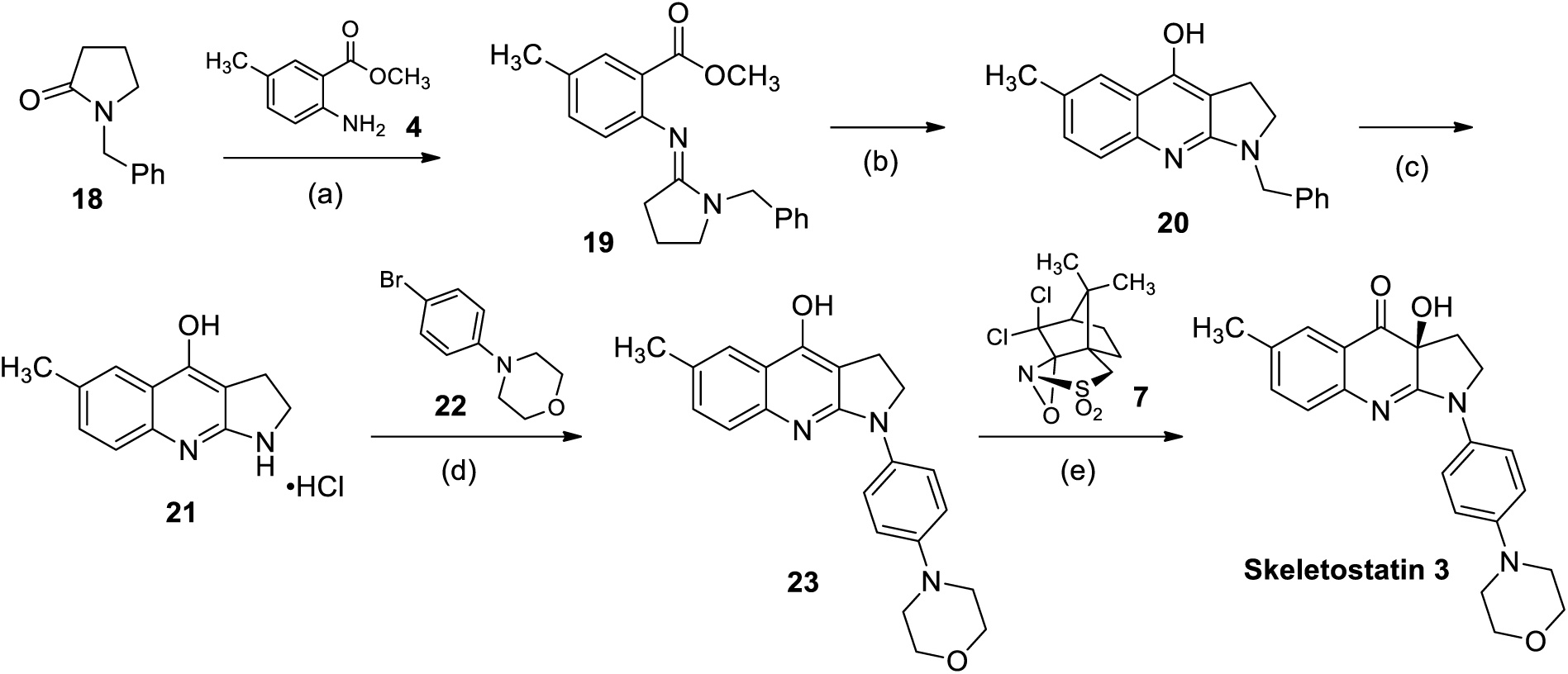
Synthesis of **Skeletostatin 3**. Reagents and conditions: (a) **4**, POCl_3_, CH_2_Cl_2_, reflux, 16 h, 81%; (b) LiHMDS, THF, 0 °C, 2 h, 42%; (c) AlCl_3_, benzene, reflux, 3 h, quantitative; (d) CuI, Cs_2_CO_3_, NaI, and *N,N*’-dimethyl-(1*R*,2*R*)-1,2-cyclohexanediamine, 1,4-dioxane, 100 °C, 64 h, 17%; (e) LiHMDS, THF, –10 °C, 2 h, 38%

### Solubility and photostability

Kinetic aqueous solubility and photostability were determined for all compounds (**Table 1**). Skeletostatin 1 and 2 showed a ∼4-fold improvement in solubility over blebb, while skeletostatin 3 showed a ∼2-fold reduction. The photostability of skeletostatins 1 and 3 was markedly improved compared to blebb after both 4 and 24 hours of bright light exposure. Skeletostatin 2 showed reduced photostability.

**Table 1.** Kinetic aqueous solubility and photostability of blebb and skeletostatins.

| Compound | Solubility (μM) | Photostability |  |  |  |
| --- | --- | --- | --- | --- | --- |
|  |  | 0 h | Light, 4 h | Light, 24 h | Dark, 24 h |
| Blebbistatin | 21 | >99.0% | 82.9% | 9.3% | >99.0% |
| Skeletostatin 1 | 78 | >99.0% | 95.4% | 62.9% | >99.0% |
| Skeletostatin 2 | 79 | 98.9% | 12.1% | 14.3% | 97.3% |
| Skeletostatin 3 | 8.3 | >99.0% | 90.8% | 43.6% | 96.2% |

### Spectral characterization of skeletostatins

First, we attempted to determine the absorbance spectra of blebb and skeletostatins in PBS containing 1% DMSO at a final concentration of 5 µM at room temperature. All compounds are soluble under these conditions (**Table 1**). Because light absorption was too weak to achieve an acceptable signal to noise ratio (data not shown), all absorbance spectra were determined in DMSO at a concentration of 0.5 mM (**Fig. 1A**). Blebb showed several overlapping peaks in the UV range and one broad absorption peak at 422 nm. Skeletostatins showed similar spectra with various levels of shifts both in the intensity and position of individual peaks (**Fig. 1A** and **Table 2**). Absorption of violet and blue light in the range of 400 to 550 nm is consistent with the yellowish color of these compounds.

**Table 2.** Molar extinction coefficients (ε) of blebb and skeletostatins at the main absorption peaks (λ). Absorbance spectra were recorded in DMSO. See Fig. 1A for further details.

| Blebbistatin |  | Skeletostatin 1 |  | Skeletostatin 2 |  | Skeletostatin 3 |  |
| --- | --- | --- | --- | --- | --- | --- | --- |
| $\lambda$ (nm) | $\epsilon$ (M<sup>-1</sup>cm<sup>-1</sup>) | $\lambda$ (nm) | $\epsilon$ (M<sup>-1</sup>cm<sup>-1</sup>) | $\lambda$ (nm) | $\epsilon$ (M<sup>-1</sup>cm<sup>-1</sup>) | $\lambda$ (nm) | $\epsilon$ (M<sup>-1</sup>cm<sup>-1</sup>) |
| 272 | 23600 | 272 | 26420 | 278 | 20560 | 278 | 17580 |
| 301 | 13840 | 300 | 11760 | 307 | 18920 | 308 | 15940 |
| 310 | 12660 | 326 | 15700 | 315 | 18940 | 313 | 15920 |
| 422 | 8540 | 413 | 12320 | 431 | 11000 | 441 | 10260 |

**Figure 1.**
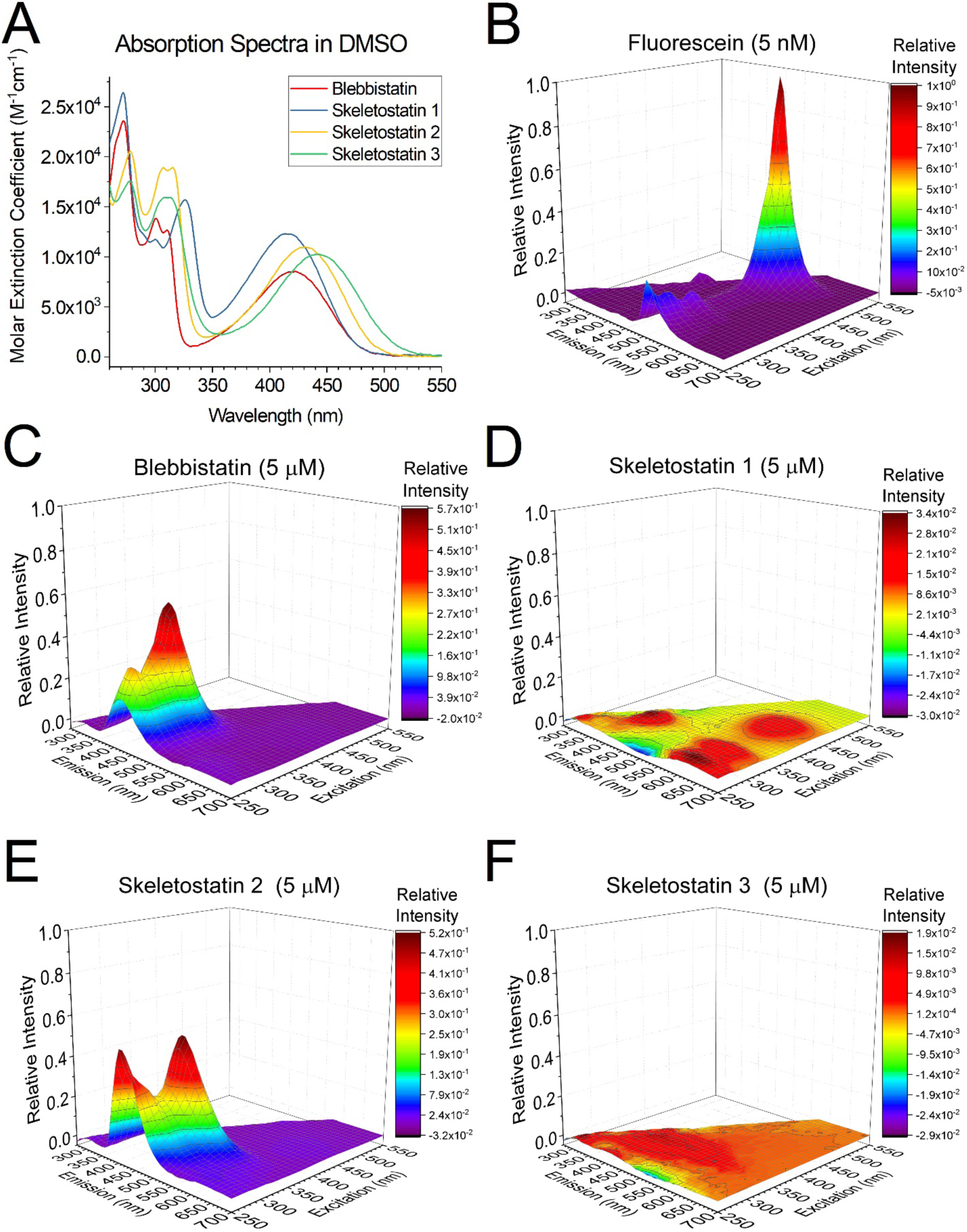
Absorption and fluorescence spectra of blebbistatin and skeletostatins. **(A)** UV-VIS absorption spectra for blebb and skeletostatins were recorded in DMSO at a concentration of 0.5 mM. Background subtracted spectra are reported as molar extinction coefficients (ε). No spectral information could be obtained at wavelengths less than 260nm due to the strong light absorption of DMSO. No compounds showed light absorption at wavelengths beyond 550 nm. Molar extinction coefficients recorded at the spectral peaks are reported in Table 2. 3D Fluorescence spectra were recorded for **(B)** fluorescein, **(C)** blebb and **(D-F)** skeletostatins 1-3 in PBS containing 1% DMSO. The concentration of fluorescein was 5 nM, while the concentration of all other compounds was 5 µM. Relative intensities are shown to facilitate comparisons. (Note that the scale is the same on all Z-axes.) Since skeletostatin 1 and 3 showed very weak fluorescence, the surfaces are also color coded as heatmaps to visualize peaks better. See Table 3 for a comparison of peak intensities and Supplementary Table 1 for spectral data.

Some studies have reported that blebb exhibits high levels of fluorescence, which has long been considered a significant issue in imaging applications *in vivo*^*28, 29, 31, 42*, [12a, 12b, 15-16]^. For example, Lucas-Lopez et al. reported that blebb shows significant emission in the green fluorescent protein (GFP) emission wavelength range with an emission maximum at 601 nm following excitation at 440 nm, which may limit its application in fluorescence-based imaging experiments involving GFP in live cells^[15]^. Blebb’s fluorescence may also interfere with some FRET-based applications^[16]^. Other studies used different excitation (λ_ex_) and emission (λ_em_) wavelengths to assess the fluorescent properties of blebb, producing largely incomparable data. Kepiro et al. observed peak emission around ∼540 nm using λ_ex_ = 430 nm^[12a]^. Varkuti et al. detected an emission peak at 410 nm using λ_ex_ = 350 nm^[12b]^. In a more recent study, Verhasselt et al. recorded emission spectra using λ_ex_ = 488 nm and λ_ex_ = 365 nm and observed emission peaks around ∼620 nm and ∼420 nm, respectively^[12c]^.

**Table 3.** Peaks detected in the 3D fluorescence spectra of fluorescein, blebb and skeletostatins. Peaks were defined as local intensity maxima. Relative intensity values were calculated as the ratio of the measured fluorescence intensity of the compound at the excitation (**λ_Ex_**) and emission (**λ_Em_**) wavelengths shown divided by the fluorescence intensity of fluorescein at **λ_Ex_** = 490 nm and **λ_Em_** = 510 nm extrapolated to the same concentration. Only peaks that are clearly distinguishable from noise are listed.

| Fluorescein |  |  | Blebbistatin |  |  | Skeletostatin 1 |  |  | Skeletostatin 2 |  |  | Skeletostatin 3 |  |  |
| --- | --- | --- | --- | --- | --- | --- | --- | --- | --- | --- | --- | --- | --- | --- |
| $\lambda_{ex}$ (nm) | $\lambda_{em}$ (nm) | Relative Intensity | $\lambda_{ex}$ (nm) | $\lambda_{em}$ (nm) | Relative Intensity | $\lambda_{ex}$ (nm) | $\lambda_{em}$ (nm) | Relative Intensity | $\lambda_{ex}$ (nm) | $\lambda_{em}$ (nm) | Relative Intensity | $\lambda_{ex}$ (nm) | $\lambda_{em}$ (nm) | Relative Intensity |
| 490 | 510 | 1 | 340 | 410 | 5.68x10<sup>-4</sup> | 330 | 400 | 3.42x10<sup>-5</sup> | 340 | 440 | 5.24x10<sup>-4</sup> | 260 | 310 | 1.93x10<sup>-5</sup> |
| 250 | 510 | 2.19x10<sup>-1</sup> | 270 | 420 | 3.22x10<sup>-4</sup> | 270 | 620 | 3.40x10<sup>-5</sup> | 250 | 410 | 5.05x10<sup>-4</sup> | 330 | 400 | 1.46x10<sup>-5</sup> |
| 280 | 510 | 1.36x10<sup>-1</sup> | 270 | 620 | 2.04x10<sup>-5</sup> | 320 | 610 | 2.17x10<sup>-5</sup> | 280 | 610 | 2.84x10<sup>-5</sup> | 250 | 400 | 1.04x10<sup>-5</sup> |
| 320 | 510 | 9.46x10<sup>-2</sup> | 260 | 310 | 1.22x10<sup>-5</sup> | 260 | 310 | 1.95x10<sup>-5</sup> | 260 | 310 | 1.60x10<sup>-5</sup> |  |  |  |
| 400 | 420 | 6.00x10<sup>-2</sup> | 440 | 630 | 9.64x10<sup>-6</sup> | 420 | 610 | 1.87x10<sup>-5</sup> | 440 | 630 | 1.27x10<sup>-5</sup> |  |  |  |

It is important to use a method that is amenable to the comprehensive characterization of the fluorescent properties of blebb and skeletostatins and that enables generation of data that is comparable across studies. To avoid precipitation-based artifacts, all compounds were dissolved in DMSO and diluted 100x in PBS to a final concentration of 5 µM. Diluted compound solutions were transferred to a cuvette and fluorescence 3D spectra were recorded over a wide range of excitation (250-550) and emission (270-700) wavelengths (**Fig. 1B-F**). Excitation was limited to the range of 250 nm to 550 nm because DMSO shows very strong light absorption at wavelengths < 250 nm and no compounds absorbed light at wavelengths > 550 nm. A solution of fluorescein at 5 nM concentration was used as a reference. As expected, fluorescein showed strong fluorescence at λ_ex_ = 490 nm, λ_em_ = 490 nm (**Fig. 1B**). The intensity detected at this peak was used in all subsequent experiments to calculate relative spectra facilitating comparisons greatly. Peak positions and relative peak intensities for all compounds are reported in **Table 3**. In agreement with the results of Verhasselt et al.^[12c]^, 5 uM blebb showed only relatively weak fluorescence (**Fig. 1C**). Even at the two highest peaks in blebb’s spectrum (λ_ex_ = 340 nm, λ_em_ = 410 nm and λ_ex_ = 270 nm, λ_em_ = 420 nm), the intensity of the emitted light was ∼1800x and ∼3100x less compared to the peak of fluorescein. Fluorescence of blebb was barely measurable in the wavelengths used to excite GFP. Supersaturated levels of blebb in a solution eventually lead to precipitation and the resulting blebb crystals do show strong fluorescence^*[12c]*^. Previously, we have also observed the strong fluorescence of blebb crystals in live cell imaging experiments using λ_ex_ = 405 nm and λ_em_ = 455 nm^[17]^. Interestingly, these crystals showed much weaker fluorescence when excited at 488 nm (λ_em_ = 525 nm). Nevertheless, the 3D fluorescence spectra presented here clearly show that blebb’s fluorescence will not interfere with fluorescence-based imaging applications at these wavelengths when the compound is in solution.

The spectrum of skeletostatin 2 is similar to blebb (**Fig. 1E** and **Table 3**; peaks at λ_ex_ = 340, λ_em_ = 440 nm and λ_ex_ = 250 nm, λ_em_ = 410 nm). Although this compound shows relatively weak fluorescence emission at these peaks (more than 3 orders of magnitude weaker than fluorescein at λ_ex_ = 490 nm, λ_em_ = 510 nm), it may interfere with certain fluorophores in several experimental setups. On the other hand, its fluorescence could potentially be exploited in other experiments (e.g. following their binding to myosins *in vitro*). Skeletostatin 1 and 3 were practically non-fluorescent (**Fig. 1D** and **F, Table 3**), which may represent a significant advantage in *in vivo* imaging applications. All 3D spectra are reported in **Supplementary Table 1**.

### Inhibition of the *in vitro* ATPase activity of class II myosins by blebbistatin and skeletostatins

The potency of blebb’s inhibitory activity against all members of the myosin II family was first benchmarked by biochemical ATPase assays^[18]^, confirming the greatest inhibition being for SkMII (K_I_ = 0.28 µM; **Fig. 2A, Table 4**). With the goal of developing a blebb derivative with improved selectivity for SkMII over CMII (K_I_ = 1.9 µM; selectivity = 6.8-fold), the inhibitory activity of all three skeletostatin compounds was next determined against these two myosin II family members. Skeletostatin 1, 2 and 3 all inhibited SkMII at submicromolar concentrations, similar to blebb (**Fig. 2B-D**). However, unlike blebb, the skeletostatins all displayed strong selectivity for SkMII over CMII (**Table 4**). Indeed, Skeletostatin 1 displayed a 173-fold selectivity for SkMII with a CMII K_I_ of 83 uM. Further, because bovine CMII (MYH7) protein was used, this also reflects selectivity for fast versus slow (also MYH7) SkMII^[19]^. Given Skeletostatin 1’s optimal physicochemical properties, potency and selectivity, its inhibition of SmMII and NMII were determined. As with CMII, Skeletostatin 1’s selectivity for SkMII over SmMII and NMII was improved relative to blebb (**Fig. 2B, Table 4**).

**Table 4.** Summary of ATPase screening data. Selectivity was defined as the ratio of the inhibitory constants (e.g. K_I,CMII_/K_I,SkMII_). A ratio over 1.0 represents selectivity for SkMII.

| Compound | ATPase Assay, $K_i$ (μM) | | | | Selectivity | | |
| --- | --- | --- | --- | --- | --- | --- | --- |
|  | SkMII | CMII | SmMII | NMII | CMII | SmMII | NMII |
| Blebbistatin | 0.28 | 1.9 | 3.2 | 2.9 | 6.8 | 11.4 | 10.4 |
| Skeletostatin 1 | 0.48 | 83 | 19 | 22 | 172.9 | 39.6 | 45.8 |
| Skeletostatin 2 | 0.27 | 34 | ND | ND | 125.9 | ND | ND |
| Skeletostatin 3 | 0.79 | >56<sup>[a]</sup> | ND | ND | >70.9<sup>[a]</sup> | ND | ND |

<sup>[a]</sup>Activity was not detected even at the highest concentration tested (solubility limited). Therefore, a lower limit for $K_i$ was estimated based on the highest concentration achieving less than 10% inhibition. ND: not determined.

**Figure 2.**
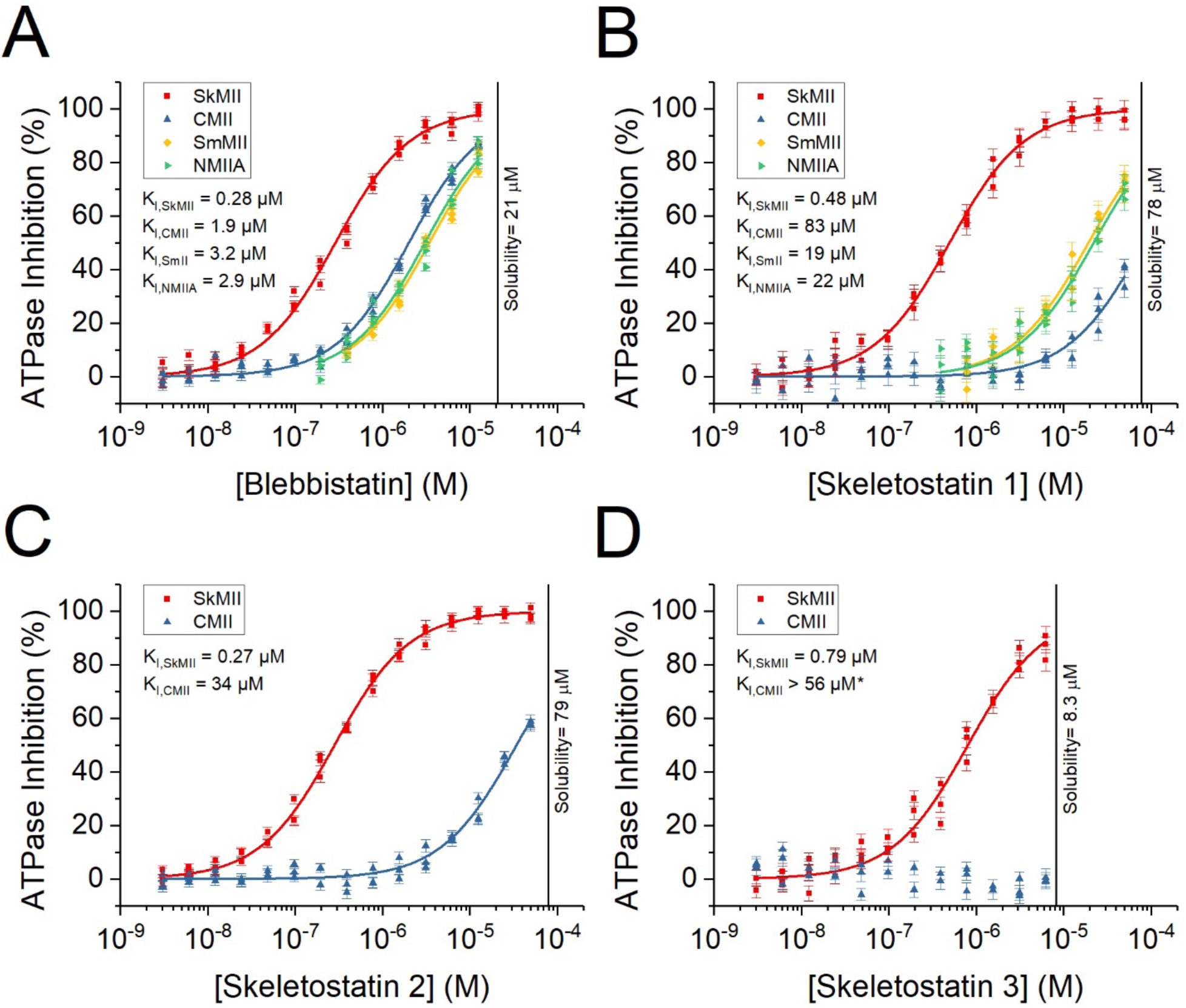
ATPase assays. Evaluation of the effects of blebbistatin (A), skeletostatin 1 (B), skeletostatin 2 (C), and skeletostatin 3 (D) on the ATPase activity of SKMII (red), CMII (blue), SmMII (yellow), and NMIIA (green) was performed in an NADH linked ATPase assay^[18]^. Inhibitory constants (K_I_) were determined by fitting the dose response data to a quadratic equation corresponding to a simple 1:1 equilibrium binding model^[18]^. Skeletostatins showed similar SkMII potency, but highly improved selectivity compared to blebb.

### Inhibition of cytokinesis by blebbistatin and skeletostatins

Blebb interferes with cytokinesis through NMII inhibition, resulting in the accumulation of multinucleated cells in cell cultures^[3]^. Therefore, we utilized cytokinesis as a cell-based assay to determine potential cytotoxicity of blebb and the skeletostatins, and to confirm cell membrane penetrance^[17]^ The results were further used to assess NMII activity. COS7 cells, which primarily express the NMIIB isoform^[20]^, were treated with a concentration series of Skeletostatin 1, 2, 3 or blebb on 96-well plates. Cytotoxicity was calculated as the ratio of dead nuclei to total nuclei^[17]^. None of the skeletostatins, nor blebb, showed any sign of cytotoxicity, even when applied at saturation levels (slightly above solubility; **Fig. 3**). The nuclei-to-cell ratio, a measure of polyploidy that reflects cytokinesis defects^[17]^, was then calculated to determine compound potencies (EC_50_). Blebb showed an EC_50_ of 1.9 µM in the cytokinesis assay (**Fig. 3A, Table 5**). Consistent with the NMII ATPase assay results, the potency of skeletostatin 1 was highly reduced (35 µM) relative to blebb. Skeletostatin 2 showed very low activity, with an EC_50_ of 96 µM. While limited by solubility, skeletostatin 3 showed a total lack of activity (EC_50_ > 6.7 µM).

**Table 5.**
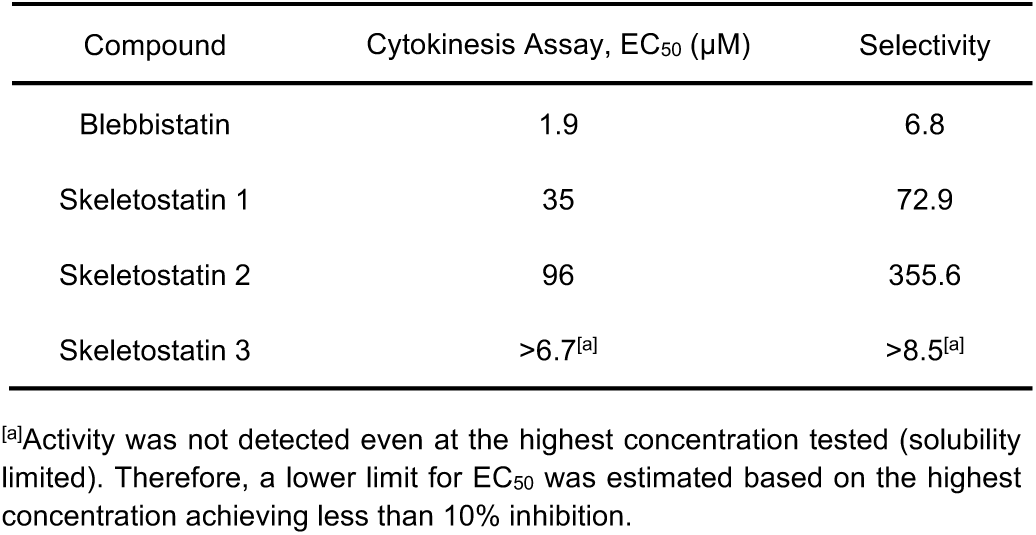
Half maximal effective concentrations (EC_50_) determined in the cytokinesis assay. Selectivity was defined as the ratio of the EC_50_ and the inhibitory constant determined in the SkMII ATPase assay (EC_50_/K_I,SkMII_). A ratio over 1.0 represents selectivity for SkMII.

**Figure 3.**
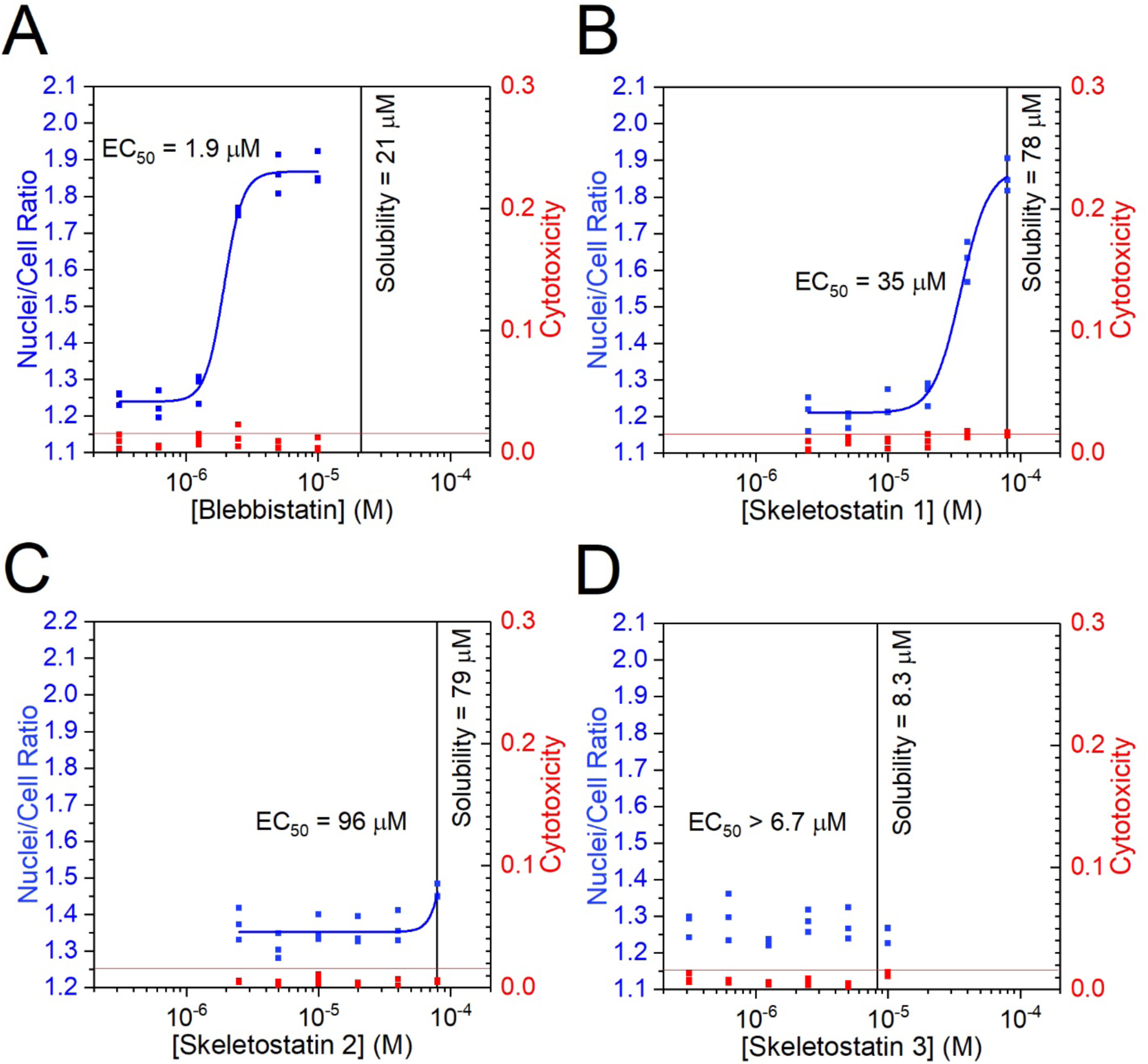
Cytokinesis assay. The activity of blebb (A) and skeletostatins (B-D) was evaluated in a cytokinesis assay as published previously^[17]^. Skeletostatins showed reduced cytokinesis inhibition compared to blebb. Each datapoint (blue) represents the nuclei-to-cell ratio (left axes) determined for a single well. Cytotoxicity was also quantified as the dead nuclei to total nuclei ratio (red, right axes). Red horizontal lines represent the empirical threshold above which cytotoxicity is considered significant^[17]^. Kinetic aqueous solubilities are represented by vertical black lines. Although the solubility in culture medium is generally higher than the kinetic solubility due to the presence of proteins, compound precipitates usually influence dose-response curves^[17]^. Therefore, highest compound concentrations tested in this assay were chosen to roughly match the reported kinetic aqueous solubilities.

### *In vivo* tolerability and efficacy profiling

Skeletostatin 1 displayed excellent solubility (78 µM), improved photostability (63% remaining after 24 hours of illumination by bright light), no cytotoxicity, submicromolar SkMII potency (0.48 µM), and selectivity heavily biased for SkMII. Based on skeletostatin 1’s weak inhibition of CMII and SmMII, we hypothesized that *in vivo* systemic administration would be well-tolerated, even at relatively high doses intended to measurably inhibit skeletal muscle function.

To obtain an initial measure of tolerability, mice were injected with vehicle or skeletostatin 1 at 2 mg/kg, 5 mg/kg, and 10mg/kg (IP) and allowed to freely explore an open field. Total distance moved did not differ between the groups at the 5 or 60 minute assessments (5 min: F_(3.10)_=1.911, p=0.1918; 60 min: F_(3,12)_=0.3409, p=0.7962; **Fig. 4**). Each mouse was visually assessed every 5 minutes during open field testing to determine any general health effects of skeletostatin 1. Two different methods were used, Body Condition Scoring (BCS; scale is 1-5, 3 being optimal) and Pain and Distress Assessment Scale (scale is 1-5, 1 indicating no pain)^[21]^. All mice were rated as maintaining a BCS 3 and a 1 on the Pain and Distress Assessment Scale throughout the observation period for all doses, indicating that at every assessment, mice remained well-conditioned, maintained grooming, were alert, active, and displayed normal eye, ear, and body posture as well as movement, suggesting excellent tolerability, even at 10 mg/kg.

**Figure 4.**
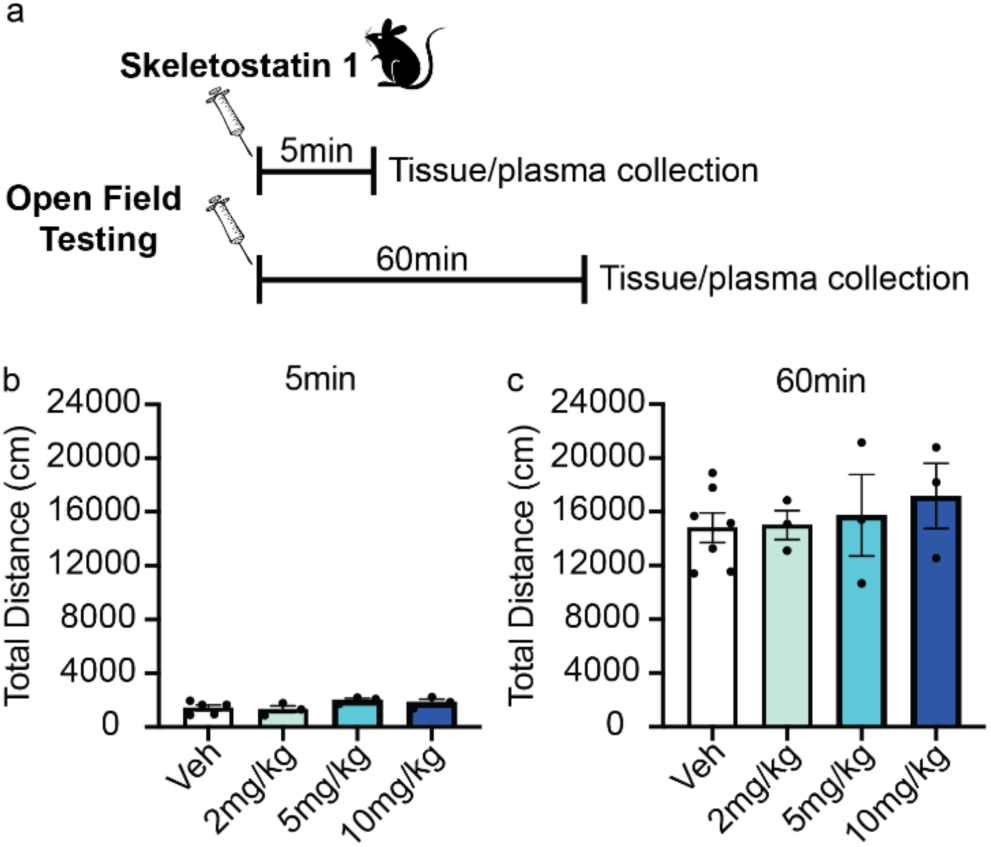
Skeletostatin 1 did not affect general locomotion in Open Field. **(A)** Schematic of behavioral methods. Mice were allowed to freely explore an open field box for **(B)** 5 or **(C)** 60min following injection of vehicle or Skeletostatin 1 (2, 5, or 10 mg/kg). No significant differences were observed between any groups. Error bars are ±SEM.

Because no gross effects on locomotion were observed at any dose of skeletostatin 1 and because 2, 5 and 10 mg/kg doses were all well-tolerated, we next assessed the effect of 5 and 10 mg/kg on coordinated motor function using the rotarod test. Rotarod was selected because it provides a more sensitive analysis of muscle and motor function than the open field^[22]^. Mice were injected with vehicle or 5 mg/kg or 10 mg/kg of skeletostatin 1 (IP) before undergoing 2 rotarod testing trials. Each trial lasted until a mouse fell off, or after 5 minutes, whichever came first, with an intertrial interval of 40 minutes. The first trial began 5 minutes after injection. Trials were analyzed separately to account for changing levels of skeletostatin 1 in muscle at the different time points (see next section). For Trial 1, there was a trend towards decreased motor coordination at both skeletostatin 1 doses (F_(2,18)_=2.464, p=0.1133; **Fig. 5**). The lack of significance was driven by the cap of 300 seconds placed on a test period and the natural variability present in control animals experiencing a rotating rod for the first time. Indeed, two vehicle animals fell off within 90 seconds of the trial start, while another vehicle animal reached the maximum trial period of 300 seconds without falling off and was removed from the apparatus. For Trial 2, which began 50 minutes after skeletostatin 1 injection and 40 minutes after the completion of Trial 1, ANOVA revealed a significant difference between groups (F_(2,18)_=6.772, p=0.0064; **Fig. 5**), and post hoc analysis confirmed that vehicle-treated mice stayed on the rotarod longer than mice treated with 10 mg/kg (p=0.0054) of skeletostatin 1. There was a strong statistical trend for reduced motor function following treatment with 5 mg/kg skeletostatin 1 relative to vehicle (p=0.0640), and mice treated with 5 and 10mg/kg were not significantly different from each other (p=0.4781). As reported during open field, no adverse clinical signs were observed in the skeletostatin 1-treated mice. Overall, these results demonstrate that skeletostatin 1 disrupts complex motor function (i.e. walking on a narrow, rotating rod) at a well-tolerated dose of 10 mg/ kg, consistent with its preferential inhibition of SkMII over the other myosin IIs.

**Figure 5.**
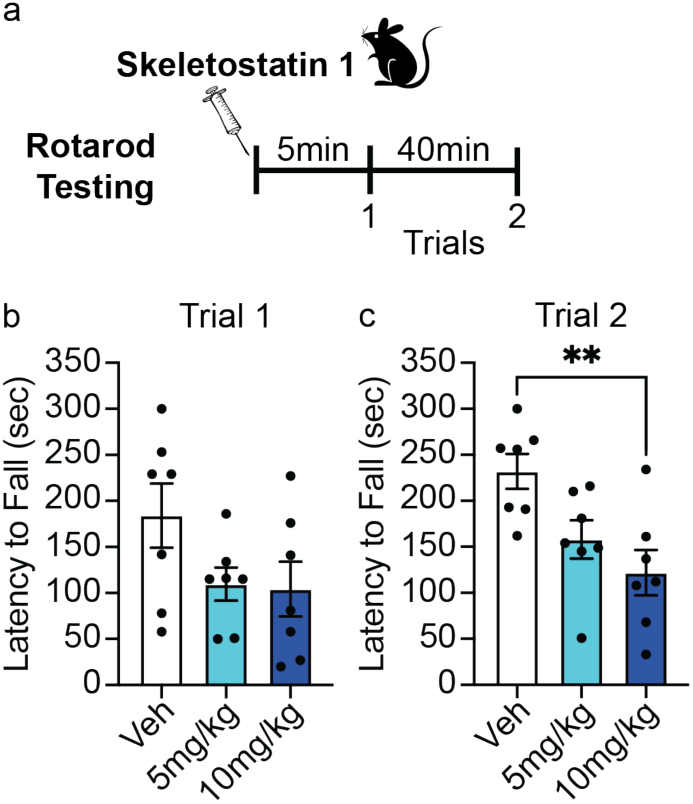
Skeletostatin 1 attenuates motor function. (a) Schematic of behavioral methods. Skeletostatin 1-treated mice (5 and 10 mg/kg) 5min prior to rotarod (b) trial 1 and 45min before (c) trial 2 had lower latencies to fall off compared to vehicle-treated mice. Error bars represent standard error of the mean and **p < 0.01.

### *In vivo* pharmacokinetic profiling

Finally, skeletostatin 1 concentrations in plasma and skeletal muscle were determined 5 and 60 minutes after injection by mass spectrometry (**Table 6**). High levels of the compound were achieved at both doses in plasma and muscle, with a consistent muscle to plasma ratio of approximately 2.8 at both time points and doses. Further, there was an approximate 10-fold reduction in skeletostatin 1 levels in both plasma and muscle by 60 minutes, a time point that corresponded to 10 minutes after Trial 2. This relatively rapid clearance indicates skeletostatin 1 is ideal when a short-acting SkMII inhibitor is desired.

**Table 6.** Pharmacokinetics. Average concentration (µM) in plasma or skeletal muscle after 5 or 60 min following IP injections of 5 mg/kg or 10 mg/kg skeletostatin 1.

|  |  | 5 min |  |  | 60 min |  |  |
| --- | --- | --- | --- | --- | --- | --- | --- |
| Dose |  | Plasma (μM) | Muscle (μM) | Muscle: Plasma | Plasma (μM) | Muscle (μM) | Muscle: Plasma |
| 5 mg/kg | AVG | 0.84 | 2.28 | 2.86 | 0.08 | 0.22 | 2.80 |
|  | SEM | 0.54 | 1.37 | 0.14 | 0.02 | 0.05 | 0.16 |
| 10 mg/kg | AVG | 2.05 | 5.68 | 2.79 | 0.19 | 0.52 | 2.76 |
|  | SEM | 0.92 | 2.52 | 0.05 | 0.03 | 0.06 | 0.10 |

## Conclusion

Here we report the synthesis and physicochemical, biochemical and *in vivo* characterization of a novel blebb derivative with selectivity for SkMII over CMII, SmMII, and NMII (NMIIA and NMIIB). Following the analogy of “blebbistatin”, which inhibits cellular blebbing, the novel inhibitors reported here are referred to as “skeletostatins”. Some skeletostatins showed improved solubility and photostability compared to blebb, and some displayed highly reduced fluorescence. They were non-cytotoxic in cell culture. The compound with the best overall *in vitro* profile, skeletostain 1, was chosen for *in vivo* profiling. Skeletostatin 1 was found to have high tolerability with systemic administration, to interfere with coordinated motor function, and to clear rapidly from plasma and skeletal muscle. Together, this supports the potential of skeletostatin 1 to be used as a research tool for selectively inhibiting skeletal muscle function both *in vitro* and *in vivo*, and as an excellent starting point for further development as a therapeutic for indications involving unwanted muscle spasms and neuromuscular disorders, such as Duchene Muscular Dystrophy^[9]^.

## Acknowledgements

This work was supported by a grant from the National Institute of Neurological Disorders and Stroke and the National Institute on Drug Abuse NS096833 (CAM, PRG and TMK).

## Entry for the Table of Contents

Insert graphic for Table of Contents here.

Insert text for Table of Contents here.

Institute and/or researcher Twitter usernames: @MillerLab3

