## Supporting Information for "Discovery of a Novel, Selective and Short-Acting Skeletal Myosin II Inhibitor"

**Abstract:** Insert abstract text here. Myosin IIs, actin-based motors that utilize the chemical energy of ATP to generate force, have potential as therapeutic targets. Their heavy chains differentiate the family into muscle (skeletal [SkMII], cardiac, smooth) and nonmuscle myosin IIs. Despite therapeutic potential for muscle disorders, no SkMII-specific inhibitor has been reported and characterized. Here we present the discovery, synthesis and characterization of “skeletostatins”, novel derivatives of the pan-myosin II inhibitor blebbistatin, with selectivity within the myosin IIs for SkMII. In addition, the skeletostatins bear improved potency, solubility and photostability, without cytotoxicity. Based on its optimal *in vitro* profile, Skeletostatin 1’s *in vivo* tolerability, efficacy and pharmacokinetics were determined. Skeletostatin 1 was well-tolerated in mice, impaired motor performance, and had an excellent muscle to plasma ratio. Skeletostatins are useful probes for basic research and a strong starting point for drug development.

### Experimental Procedures

#### 1. Chemistry

#### 1.1. General Information

Unless otherwise stated, commercially available reagents and solvents were used without purification. All reactions were performed under an atmosphere of nitrogen, unless otherwise noted. Temperatures provided are external temperatures of the heating or cooling bath, unless otherwise noted. Flash column chromatography was performed on an automated Biotage Isolera flash chromatography system, using Biotage SNAP Ultra silica cartridges. Melting points were determined on a Mel-Temp apparatus and are uncorrected. NMR spectra were recorded on a Bruker spectrometer (300 or 500 MHz) with chemical shifts ($\delta$) reported in parts-per-million (ppm) relative to tetramethylsilane or the residual signal of the deuterated solvent. $^1$H NMR data are reported as follows: chemical shift, multiplicity (s = singlet, d = doublet, t = triplet, q = quartet, m = multiplet, and br = broad), coupling constant in Hz, and number of protons. Mass spectra were obtained using a Waters LC/MS system using electrospray ionization. Purity was determined by HPLC analysis on a Varian system using an xBridge 3.5 µm C18 column (150 × 4.6 mm) or a YMC ODS-AQ C18 120 Å column (150 × 4.6 mm) using a gradient of acetonitrile in water containing 0.1% trifluoroacetic acid (see HPLC analytical methods for details). Peaks were detected at a wavelength of 254 nm with a photodiode array detector. The chiral purity was determined via chiral HPLC analysis using a Chiralpak AD 5 µm column (250 × 4.6 mm) or a Chiralpak OD 5 µm column (250 × 4.6 mm) using a gradient of *i*-propyl alcohol in heptane (see HPLC analytical methods for details). Detection wavelength was set at 254 nm or 270 nm. Analyses were performed at ambient temperature (20–25 °C).

#### 1.2. HPLC Conditions:

**Method A**
Column: xBridge 3.5 µm C18 (150 × 4.6 mm)
Mobile Phase A: Water containing 0.1% v/v Trifluoroacetic Acid
Mobile Phase B: Acetonitrile containing 0.1% v/v Trifluoroacetic Acid
Detection: 254 nm

**Method A Gradient**

| Time (min) | Flow (mL/min) | %A | %B |
|---|---|---|---|
| 0.0 | 1.0 | 95.0 | 5.0 |
| 20.0 | 1.0 | 0.0 | 100.0 |
| 25.0 | 1.0 | 0.0 | 100.0 |

**Method B**
Column: YMC ODS-AQ C18 120 Å (150 × 4.6 mm)
Mobile Phase A: Water containing 0.1% v/v Trifluoroacetic Acid
Mobile Phase B: Acetonitrile containing 0.1% v/v Trifluoroacetic Acid
Detection: 254 nm

**Method B Gradient**

| Time (min) | Flow (mL/min) | %A | %B |
|---|---|---|---|
| 0.0 | 1.0 | 95.0 | 5.0 |
| 15.0 | 1.0 | 0.0 | 100.0 |
| 19.0 | 1.0 | 0.0 | 100.0 |

#### 1.3. Chiral HPLC Conditions:

**Method A**
Column: Chiralpak AD 5 µm (250 × 4.6 mm)
Mobile Phase A: Heptane
Mobile Phase B: *i*-Propyl Alcohol
Detection: 254 nm

**Method A Gradient**

| Time (min) | Flow (mL/min) | %A | %B |
|---|---|---|---|
| 0.0 | 1.0 | 90.0 | 10.0 |
| 5.0 | 1.0 | 90.0 | 10.0 |
| 20.0 | 1.0 | 50.0 | 50.0 |
| 35.0 | 1.0 | 50.0 | 50.0 |

**Method B**

Column: Chiralpak OD 5 µm (250 × 4.6 mm)
Mobile Phase A: Heptane
Mobile Phase B: *i*-Propyl Alcohol
Detection: 270 nm

**Method B Gradient**

| Time (min) | Flow (mL/min) | %A | %B |
|---|---|---|---|
| 0.0 | 1.0 | 90.0 | 10.0 |
| 2.0 | 1.0 | 90.0 | 10.0 |
| 18.0 | 1.0 | 50.0 | 50.0 |
| 35.0 | 1.0 | 50.0 | 50.0 |

#### 1.4. Preparation of (S)-3a-Hydroxy-6-methyl-1-phenyl-3,3a-dihydro-1H-pyrrolo[2,3-b]quinolin-4(2H)-one (Blebbistatin)[1]

**Synthetic Scheme**

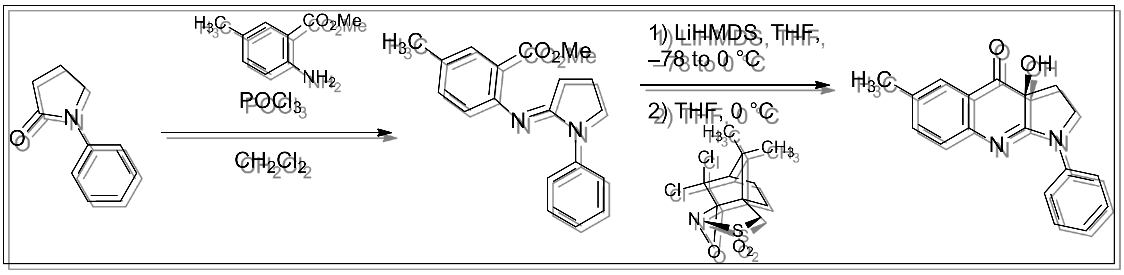

**Preparation of Methyl 5-Methyl-2-((1-phenylpyrrolidin-2-ylidene)amino)benzoate**

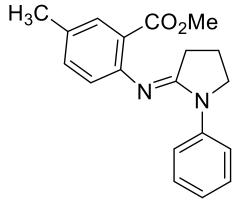

A solution of 1-phenylpyrrolidin-2-one (2.00 g, 12.4 mmol) in methylene chloride (25 mL) was treated with phosphorous oxychloride (1.8 mL, 3.0 g, 19 mmol) and stirred under a nitrogen atmosphere at ambient temperature for 4 h. The mixture was treated with a solution of methyl 2-amino-5-methylbenzoate (2.06 g, 12.5 mmol) in methylene chloride (10. mL) and heated at 45 °C for 40.5 h. After this time, the reaction mixture was allowed to cool to ambient temperature and was treated with sodium bicarbonate (20. mL). The organic and aqueous layers were separated, and the aqueous layer was washed with ethyl acetate (2 × 20 mL). The combined organics were dried over sodium sulfate, filtered, and concentrated under reduced pressure. The residue was dissolved in ethyl acetate (20 mL) and extracted with 0.3 M hydrochloric acid (4 × 20 mL). The combined acid layers were adjusted to pH ~11 with 2.0 M aqueous sodium hydroxide and extracted with ethyl acetate (3 × 50 mL). The combined organics were dried over sodium sulfate, filtered, and concentrated under reduced pressure to provide methyl 5-methyl-2-((1-phenylpyrrolidin-2-ylidene)amino)benzoate (2.06 g, 54%) as a pale yellow oil: $^1$H NMR (300 MHz, DMSO-$d_6$) δ 7.83 (d, *J* = 8.1 Hz, 2H), 7.54 (s, 1H), 7.35 (apparent t, *J* = 7.2 Hz, 2H), 7.26 (d, *J* = 8.7 Hz, 1H), 7.09–7.02 (m, 1H), 6.76–6.67 (m, 1H), 3.85 (t, *J* = 6.0 Hz, 2H), 3.72 (s, 3H), 2.44–2.36 (m, 2H), 2.28 (s, 3H), 2.03–1.93 (m, 2H); ESI MS *m/z* 309 [$C_{19}H_{20}N_2O_2$+ H]$^+$.

**Preparation of (*S*)-3a-Hydroxy-6-methyl-1-phenyl-3,3a-dihydro-1*H*-pyrrolo[2,3-*b*]quinolin-4(2*H*)-one**

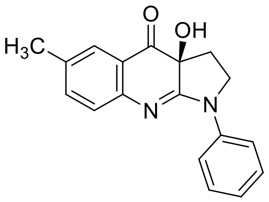

A solution of methyl 5-methyl-2-((1-phenylpyrrolidin-2-ylidene)amino)benzoate (2.06 g, 6.68 mmol) in tetrahydrofuran (20 mL) was cooled in a dry ice/acetone bath under a nitrogen atmosphere and treated dropwise with a 1.0 M solution of lithium bis(trimethylsilyl)amide in tetrahydrofuran (20.0 mL, 20.0 mmol). The mixture was stirred for 45 min, during which time the bath temperature increased to ~0 °C. After 45 min, the mixture was treated with a solution of (-)-(8,8-dichlorocamphorylsulfonyl)oxaziridine (3.99 g, 13.4 mmol) in tetrahydrofuran (10 mL) and was stirred for 40 min. After this time, the mixture was treated with saturated aqueous ammonium iodide (6 mL) followed by saturated aqueous sodium thiosulfate (12 ml) and brine (30 mL). The organic and aqueous layers were separated, and the aqueous layer was washed with ethyl acetate (2 × 30 mL). The combined organics were dried over sodium sulfate, filtered, and concentrated under reduced pressure. The residue was dissolved in ethyl acetate (40 mL) and extracted with 0.3 M hydrochloric acid (4 × 40 mL). The combined acid layers were adjusted to pH ~8 with 2.0 M aqueous sodium hydroxide and extracted with ethyl acetate (4 × 70 mL). The organics were dried over sodium sulfate, filtered, and concentrated under reduced pressure. The crude product was recrystallized from hot acetonitrile to provide (*S*)-3a-hydroxy-6-methyl-1-phenyl-3,3a-dihydro-1*H*-pyrrolo[2,3-*b*]quinolin-4(2*H*)-one (898 mg, 46%) as a yellow solid: mp = 205–206 °C; $^1$H NMR (300 MHz, DMSO-$d_6$) δ 8.08 (dd, *J* = 8.7, 0.9 Hz, 2H), 7.53 (d, *J* = 1.5 Hz,1H), 7.45–7.36 (m, 3H), 7.16–7.11 (m, 2H), 6.84 (s, 1H), 4.11–4.02 (m, 1H), 3.99–3.93 (m, 1H), 2.30 (s, 3H), 2.29–2.24 (m, 2H); ESI MS *m/z* 293 [$C_{18}H_{16}N_2O_2$ + H]$^+$; HPLC (Method A) >99% (AUC), $t_R$ = 11.50 min; Chiral HPLC (Chiralpak AD, Method A) >99% (AUC), $t_R$ = 14.19 min.

#### 1.5. Preparation of (S)-3a-Hydroxy-6-methyl-1-(2-methyloxazolo[4,5-b]pyridin-6-yl)-1,2,3,3a-tetrahydro-4H-pyrrolo[2,3-b]quinolin-4-one (Skeletostatin 1)[2]

**Synthetic Scheme**

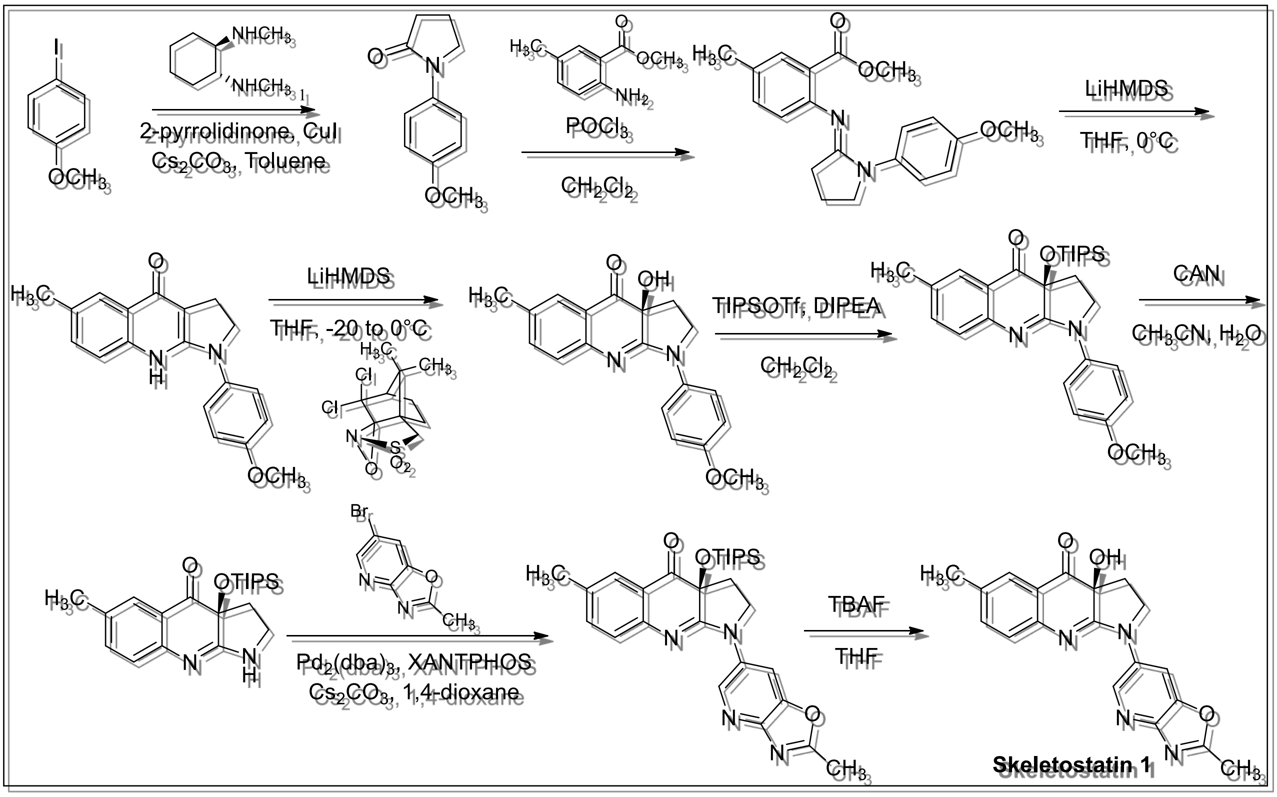

**Preparation of 1-(4-Methoxyphenyl)pyrrolidin-2-one**

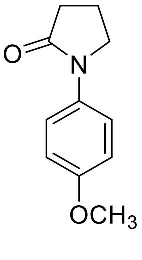

A solution of 1-iodo-4-methoxybenzene (20.0 g, 85.5 mmol) in toluene (130 mL) was treated with 2-pyrrolidinone (6.80 g, 79.9 mmol), copper iodide (1.6 g, 8.4 mmol), cesium carbonate (60.0 g, 184 mmol) and *N*,*N'*-dimethyl-(1*R*,2*R*)-1,2-cyclohexanediamine (2.30 g, 16.2 mmol) and heated at reflux under a nitrogen atmosphere for 16 h. After this time, the reaction mixture was allowed to cool to ambient temperature, diluted with ethyl acetate, and filtered through diatomaceous earth. The filtrate was concentrated under reduced pressure. The crude residue was purified by column chromatography (silica gel, 0–10% methanol/methylene chloride) to provide 1-(4-methoxyphenyl)pyrrolidin-2-one (13.2 g, 86%): $^1$H NMR (300 MHz, $CDCl_3$) δ 7.52–7.47 (m, 2H), 6.93–6.88 (m, 2H), 3.86–3.80 (m, 5H), 2.60 (t, *J* = 7.8 Hz, 2H), 2.21–2.11 (m, 2H).

**Preparation of 1-(4-Methoxyphenyl)-6-methyl-2,3-dihydro-1*H*-pyrrolo[2,3-b]quinolin-4(9*H*)-one**

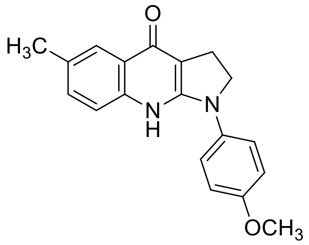

A solution of 1-(4-methoxyphenyl)pyrrolidin-2-one (10.0 g, 52.3 mmol) in dichloromethane (220 mL) was treated with phosphorous oxychloride (7.2 mL, 79 mmol) and stirred at ambient temperature for 4 h. The mixture was treated with a solution of methyl 2-amino-5-methylbenzoate (12.0 g, 72.6 mmol) in dichloromethane (40 mL) and heated at reflux for 16 h. After this time, the reaction mixture was allowed to cool to ambient temperature and concentrated under reduced pressure to remove the volatiles. The resulting residue was dissolved in ethyl acetate and washed with saturated aqueous sodium bicarbonate and water. The organic layer was extracted with 0.8 M aqueous hydrochloric acid. The combined aqueous layers were adjusted to pH ~10 with sodium hydroxide and extracted with ethyl acetate. The organics were dried over sodium sulfate, filtered, and concentrated under reduced pressure to provide methyl (*E*)-2-((1-(4-methoxyphenyl)pyrrolidin-2-ylidene)amino)-5-methylbenzoate (10.7 g, 60%). A portion of methyl (*E*)-2-((1-(4-methoxyphenyl)pyrrolidin-2-ylidene)amino)-5-methylbenzoate (10.7 g, 31.6 mmol) in tetrahydrofuran (170 mL) was cooled in a wet ice/water bath under a nitrogen atmosphere and treated with a 1 M solution of lithium bis(trimethylsilyl)amide in tetrahydrofuran (60 mL, 60 mmol), and the mixture was stirred for 2 h. After this time, the mixture was treated with water (50 mL) and saturated aqueous ammonium chloride (150 mL) and stirred at 0°C for 40 min. At this time, a precipitate formed which was isolated by filtration, washed with diethyl ether to provide 1-(4-methoxyphenyl)-6-methyl-2,3-dihydro-1*H*-pyrrolo[2,3-*b*]quinolin-4(9*H*)-one (5.7 g, 59%): ESI MS *m/z* 307 $[C_{19}H_{18}N_2O_2 + H]^+$.

**Preparation of (*S*)-3a-Hydroxy-1-(4-methoxyphenyl)-6-methyl-1,2,3,3a-tetrahydro-4*H*-pyrrolo[2,3-*b*]quinolin-4-one**

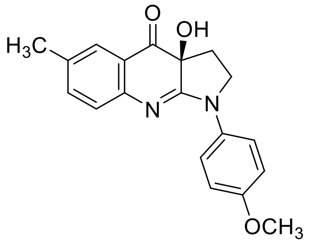

A solution of 1-(4-methoxyphenyl)-6-methyl-2,3-dihydro-1*H*-pyrrolo[2,3-*b*]quinolin-4(9*H*)-one (5.70 g, 18.6 mmol) in tetrahydrofuran (80 mL) was cooled to ~ –20°C under a nitrogen atmosphere and treated with a 1 M solution of lithium bis(trimethylsilyl)amide in tetrahydrofuran (23 mL, 23 mmol), and the mixture was stirred for 5 min. After this time, the mixture was treated with a solution of (-)-(8,8-dichlorocamphorylsulfonyl)oxaziridine (11.0 g, 36.9 mmol) and stirred for 1 h at ~ –20°C. At this time, a precipitate formed which was isolated by filtration. The filtrate was treated sequentially at 0°C with saturated aqueous ammonium iodide and saturated aqueous sodium thiosulfate and extracted with ethyl acetate. The organics were extracted with 0.8 M aqueous hydrochloric acid. The aqueous layer was basified with sodium hydroxide and extracted with ethyl acetate. The organic layers were dried over sodium sulfate, filtered, combined with the precipitate and concentrated under reduced pressure to provide (*S*)-3a-hydroxy-1-(4-methoxyphenyl)-6-methyl-1,2,3,3a-tetrahydro-4*H*-pyrrolo[2,3-*b*]quinolin-4-one (5.8 g, 96%): $^1$H NMR (500 MHz, DMSO-$d_6$) δ 8.31 (br s, 1H), 7.98–7.94 (m, 2H), 7.50 (d, *J* = 1.5 Hz, 1H), 7.34 (dd, *J* = 8.0, 2.0 Hz, 1H), 7.06 (d, *J* = 8.0 Hz, 1H), 7.01–6.98 (m, 2H), 4.07–4.02 (m, 1H), 3.92–3.88 (m, 1H), 3.77 (s, 3H), 2.29 (s, 3H), 2.26–2.23 (m, 2H); chiral HPLC (Method A) 74.2% (AUC), $t_R$ = 21.74 min.

**Preparation of (*S*)-1-(4-Methoxyphenyl)-6-methyl-3a-((triisopropylsilyl)oxy)-1,2,3,3a-tetrahydro-4*H*-pyrrolo[2,3-*b*]quinolin-4-one**

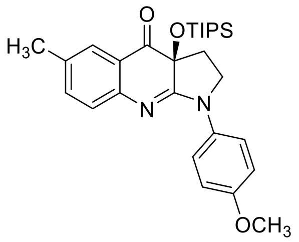

A solution of (*S*)-3a-hydroxy-1-(4-methoxyphenyl)-6-methyl-1,2,3,3a-tetrahydro-4*H*-pyrrolo[2,3-*b*]quinolin-4-one (1.70 g, 5.43 mmol) in dichloromethane (70 mL) was treated with *N*,*N*-diisopropylethylamine (3.60 mL, 20.7 mmol) and triisopropylsilyl trifluoromethanesulfonate (4.40 mL, 16.4 mmol) and heated at reflux under a nitrogen atmosphere for 16 h. After this time, the reaction mixture was allowed to cool to ambient temperature. The mixture was treated with saturated aqueous ammonium chloride. The organics were dried over sodium sulfate, filtered, and concentrated under reduced pressure. The crude residue was purified by column chromatography (silica gel, 0–60% ethyl acetate/heptane) to provide (*S*)-1-(4-methoxyphenyl)-6-methyl-3a-((triisopropylsilyl)oxy)-1,2,3,3a-tetrahydro-4*H*-pyrrolo[2,3-*b*]quinolin-4-one (1.12 g, 43%): ESI MS *m/z* 479 [$C_{28}H_{38}N_2O_3Si$ + H]$^+$.

**Preparation of (*S*)-6-Methyl-3a-((triisopropylsilyl)oxy)-1,2,3,3a-tetrahydro-4*H*-pyrrolo[2,3-*b*]quinolin-4-one**

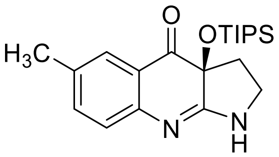

A solution of (*S*)-1-(4-methoxyphenyl)-6-methyl-3a-((triisopropylsilyl)oxy)-1,2,3,3a-tetrahydro-4*H*-pyrrolo[2,3-*b*]quinolin-4-one (1.10 g, 2.30 mmol) in acetonitrile (15 mL) was cooled in a wet ice/water bath under a nitrogen atmosphere and treated dropwise with a solution of ammonium cerium(IV) nitrate (4.00 g, 7.30 mmol) in deionized water (15 mL) and stirred for 2 h at ~0 °C. After this time, the mixture was treated with sodium thiosulfate pentahydrate (15 g) in deionized water and stirred for 30 min, followed by saturated aqueous sodium bicarbonate (150 mL) to form a slurry. The solid was removed by filtration through diatomaceous earth and rinsed with ethyl acetate. The filtrate was washed with saturated sodium bicarbonate and brine. The organics were dried over sodium sulfate, filtered, and concentrated under reduced pressure to provide (*S*)-6-methyl-3a-((triisopropylsilyl)oxy)-1,2,3,3a-tetrahydro-4*H*-pyrrolo[2,3-*b*]quinolin-4-one (850 mg, 99%): ESI MS *m/z* 373 [$C_{21}H_{32}N_2O_2Si$ + H]$^+$.

**Preparation of (*S*)-6-Methyl-1-(2-methyloxazolo[4,5-*b*]pyridin-6-yl)-3a-((triisopropylsilyl)oxy)-1,2,3,3a-tetrahydro-4*H*-pyrrolo[2,3-*b*]quinolin-4-one**

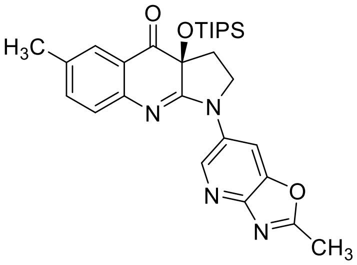

A solution of (*S*)-6-methyl-3a-((triisopropylsilyl)oxy)-1,2,3,3a-tetrahydro-4*H*-pyrrolo[2,3-*b*]quinolin-4-one (200 mg, 0.540 mmol) in 1,4-dioxane (10 mL) was treated with 6-bromo-2-methyloxazolo[4,5-*b*]pyridine (230 mg, 1.08 mmol), cesium carbonate (380 mg, 1.16 mmol), tris(dibenzylideneacetone)dipalladium(0) (70 mg, 0.076 mmol) and 4,5-bis(diphenylphosphino)-9,9-dimethylxanthene (130 mg, 0.22 mmol), purged with argon, and heated at 120 °C in a sealed vial for 16 h. After this time, the reaction mixture was allowed to cool to ambient temperature. The solids were removed by filtration through diatomaceous earth and washed with ethyl acetate. The filtrate was concentrated under reduced pressure. The crude residue was purified by column chromatography (silica gel, 0-5% methanol/methylene chloride) to provide (*S*)-6-methyl-1-(2-methyloxazolo[4,5-*b*]pyridin-6-yl)-3a-((triisopropylsilyl)oxy)-1,2,3,3a-tetrahydro-4*H*-pyrrolo[2,3-*b*]quinolin-4-one (192 mg, 71%): ESI MS *m/z* 505 [$C_{28}H_{36}N_4O_3Si$ + H]$^+$.

**Preparation of provide (*S*)-3a-Hydroxy-6-methyl-1-(2-methyloxazolo[4,5-*b*]pyridin-6-yl)-1,2,3,3a-tetrahydro-4*H*-pyrrolo[2,3-*b*]quinolin-4-one (Skeletostatin 1; BPN-0027377)**

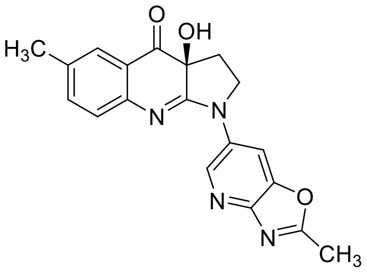

A solution of (*S*)-6-methyl-1-(2-methyloxazolo[4,5-*b*]pyridin-6-yl)-3a-((triisopropylsilyl)oxy)-1,2,3,3a-tetrahydro-4*H*-pyrrolo[2,3-*b*]quinolin-4-one (180 mg, 0.357 mmol) in tetrahydrofuran (8 mL) under a nitrogen atmosphere was treated with a 1.0 M solution of tetrabutylammonium fluoride in tetrahydrofuran (0.4 mL, 0.4 mmol), and the mixture was stirred for 1 h. After this time, the mixture was diluted with ethyl acetate and washed with water. The organic layer was dried over sodium sulfate, filtered and concentrated under reduced pressure. The crude residue was purified by column chromatography (silica gel, 0–30% (methylene chloride/methanol/ammonium hydroxide 80/18/2)/methylene chloride). The resulting residue was triturated in acetonitrile and isolated by filtration to provide (*S*)-3a-hydroxy-6-methyl-1-(2-methyloxazolo[4,5-*b*]pyridin-6-yl)-1,2,3,3a-tetrahydro-4*H*-pyrrolo[2,3-*b*]quinolin-4-one (30 mg, 24%) as a yellow solid: mp = 258–260 °C; $^1$H NMR (500 MHz, DMSO-$d_6$) δ 9.06 (dd, *J* = 8.0, 5.5 Hz, 2H), 7.56 (d, *J* = 1.5 Hz, 1H), 7.41 (dd, *J* = 8.0, 6.0 Hz, 1H), 7.20 (d, *J* = 8.0 Hz, 1H), 6.92 (s, 1H), 4.19–4.14 (m, 1H), 4.09–4.06 (m, 1H), 2.69 (s, 3H), 2.35–2.30 (m, 5H); ESI MS *m/z* 349 [$C_{19}H_{16}N_4O_3$ + H]$^+$; HPLC (Method A) >99% (AUC), $t_R$ = 11.56 min; Chiral HPLC (Chiralpak OD, Method B) 84.2% (AUC), $t_R$ = 25.04 min.

#### 1.6. Preparation of (S)-N-(4-(3a-Hydroxy-6-methyl-4-oxo-2,3,3a,4-tetrahydro-1H-pyrrolo[2,3-b]quinolin-1-yl)phenyl)acetamide (Skeletostatin 2)[3]

**Synthetic Scheme**

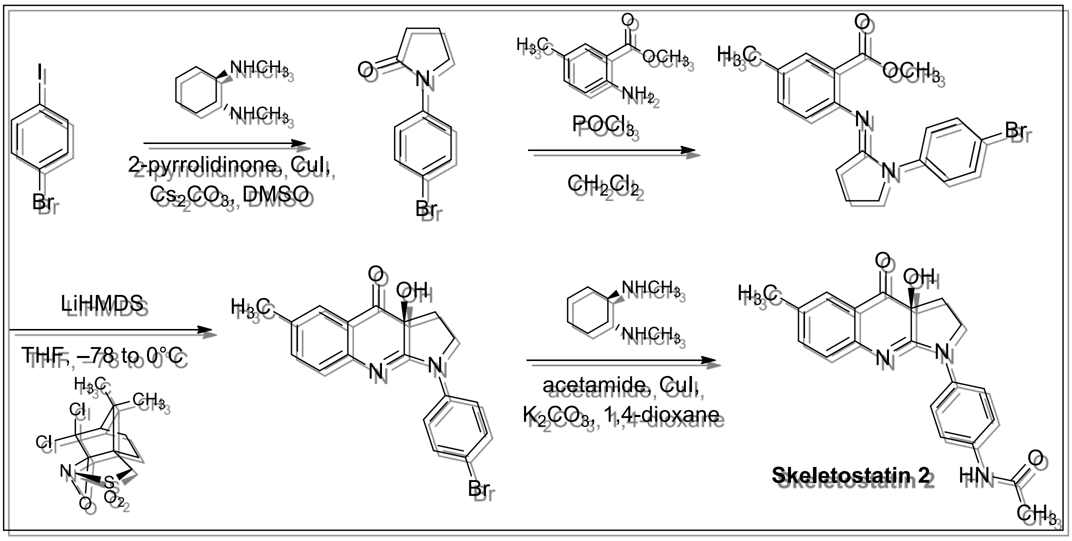

**Preparation of 1-(4-Bromophenyl)pyrrolidin-2-one**

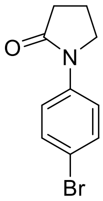

A solution of 1-bromo-4-iodobenzene (1.00 g, 3.53 mmol) in dimethyl sulfoxide (5 mL) was treated with 2-pyrrolidinone (269 μL, 3.53 mmol), copper iodide (67.0 mg, 0.353 mmol), cesium carbonate (3.46 g, 10.6 mmol) and *N,N'*-dimethyl-(1*R*,2*R*)-1,2-cyclohexanediamine (111 μL, 0.707 mmol) and heated at 110 °C under a nitrogen atmosphere for 16 h. After this time, the reaction mixture was allowed to cool to ambient temperature, diluted with water (50 mL), and extracted with ethyl acetate (3 x 50 mL). The combined organics were washed with water (4 x 5 mL), dried over sodium sulfate, filtered, and concentrated under reduced pressure. The crude residue was purified by column chromatography (silica gel, 0–60% ethyl acetate/heptane) to provide 1-(4-bromophenyl)pyrrolidin-2-one (639 mg, 75%) as a white solid: $^1$H NMR (500 MHz, $CDCl_3$) δ 7.53 (dd, *J* = 7.0, 2.5 Hz, 2H), 7.47 (dd, *J* = 7.0, 2.5 Hz, 2H), 3.84 (t, *J* = 7.0 Hz, 2H), 2.61 (t, *J* = 8.0 Hz, 2H), 2.19–2.16 (m, 2H).

**Preparation of Methyl 2-((1-(4-Bromophenyl)pyrrolidin-2-ylidene)amino)-5-methylbenzoate**

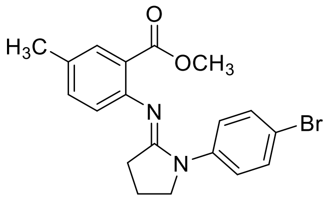

A solution of 1-(4-bromophenyl)pyrrolidin-2-one (639 mg, 2.66 mmol) in methylene chloride (25 mL) was treated with phosphorous oxychloride (0.372 mL, 3.99 mmol) and stirred under a nitrogen atmosphere at ambient temperature for 16 h. The mixture was treated with a solution of methyl 2-amino-5-methylbenzoate (440 mg, 2.66 mmol) in methylene chloride (10 mL) and heated at 45 °C for 4 d. After this time, the reaction mixture was allowed to cool to ambient temperature, quenched with saturated aqueous sodium bicarbonate (25 mL), and extracted with ethyl acetate. The organics were dried over sodium sulfate, filtered, and concentrated under reduced pressure. The crude residue was dissolved in ethyl acetate and extracted with 0.3 M hydrochloric acid (2 × 20 mL). The combined acid layers were adjusted to pH ~11 with 2.0 M aqueous sodium hydroxide and extracted with ethyl acetate (3 × 25 mL). The organics were dried over sodium sulfate, filtered, and concentrated under reduced pressure to provide methyl 2-((1-(4-bromophenyl)pyrrolidin-2-ylidene)amino)-5-methylbenzoate (396 mg, 38%) as a yellow gum: ESI MS *m/z* 389 $[C_{19}H_{19}BrN_2O_2 + H]^+$.

**Preparation of (*S*)-1-(4-Bromophenyl)-3a-hydroxy-6-methyl-1,2,3,3a-tetrahydro-4*H*-pyrrolo[2,3-*b*]quinolin-4-one**

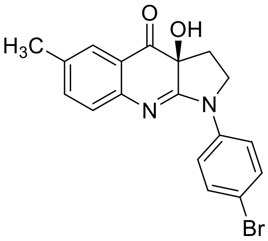

A solution of methyl 2-((1-(4-bromophenyl)pyrrolidin-2-ylidene)amino)-5-methylbenzoate (396 mg, 1.02 mmol) in tetrahydrofuran (16 mL) was cooled in a dry ice/acetone bath under a nitrogen atmosphere and treated with a 1.0 M solution of lithium bis(trimethylsilyl)amide in tetrahydrofuran (3.07 mL, 3.07 mmol). The acetone bath was replaced by a wet ice/water bath and the mixture was stirred for 1 h. After this time, the mixture was treated with a solution of (-)-(8,8-dichlorocamphorylsulfonyl)oxaziridine (762 mg, 2.56 mmol) in tetrahydrofuran (7 mL). The mixture was stirred for 2 h at ~0 °C. After this time, the mixture was treated with saturated aqueous ammonium iodide (1.2 mL) followed by saturated aqueous sodium thiosulfate (3.4 ml) followed by brine (25 mL). The aqueous layer was extracted with ethyl acetate (3 × 25 mL). The organics were combined, dried over sodium sulfate, filtered, and concentrated under reduced pressure. The crude residue was purified by column chromatography (silica gel, 12–100% ethyl acetate/heptane) and recrystallization from hot acetonitrile to provide (*S*)-1-(4-bromophenyl)-3a-hydroxy-6-methyl-1,2,3,3a-tetrahydro-4*H*-pyrrolo[2,3-*b*]quinolin-4-one (135 mg, 36%) as an orange solid: mp = 218–219 °C; $^1$H NMR (500 MHz, DMSO-$d_6$) δ 8.07 (dd, *J* = 7.0, 2.0 Hz, 2H), 7.60 (dd, *J* = 7.0, 2.0 Hz, 2H), 7.53 (apparent d, *J* = 2.0 Hz, 1H), 7.40–7.38 (m, 1H), 7.13 (d, *J* = 8.5 Hz, 1H), 6.85 (s, 1H), 4.04–4.02 (m, 1H), 3.96–3.94 (m, 1H), 2.31 (s, 3H), 2.25 (apparent d, *J* = 7.0 Hz, 2H); ESI MS *m/z* 373 $[C_{18}H_{15}BrN_2O_2 + H]^+$; HPLC (Method B) >99% (AUC), $t_R$ = 9.76 min; Chiral HPLC (Chiralpak AD, Method A) >99% (AUC), $t_R$ = 16.98 min.

**Preparation of (*S*)-*N*-(4-(3a-Hydroxy-6-methyl-4-oxo-2,3,3a,4-tetrahydro-1*H*-pyrrolo[2,3-*b*]quinolin-1-yl)phenyl)acetamide (Skeletostatin 2)**

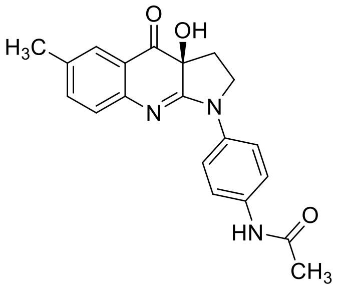

A solution of (*S*)-1-(4-bromophenyl)-3a-hydroxy-6-methyl-1,2,3,3a-tetrahydro-4*H*-pyrrolo[2,3-*b*]quinolin-4-one (50.0 mg, 0.135 mmol) in 1,4-dioxane (5 mL) was treated with acetamide (11.9 mg, 0.202 mmol), copper iodide (2.6 mg, 0.014 mmol), potassium carbonate (56 mg, 0.41 mmol) and *N,N'*-dimethyl-(1*R*,2*R*)-1,2-cyclohexanediamine (4.3 µL, 0.027 mmol) and heated at 100 °C under a nitrogen atmosphere for 16 h. After this time, the reaction mixture was allowed to cool to ambient temperature, diluted with water, and extracted with ethyl acetate (3x). The combined organics were dried over sodium sulfate, filtered, and concentrated under reduced pressure. The crude residue was purified by column chromatography (silica gel, 10–100% ethyl acetate/heptane) to provide (*S*)-*N*-(4-(3a-hydroxy-6-

methyl-4-oxo-2,3,3a,4-tetrahydro-1*H*-pyrrolo[2,3-*b*]quinolin-1-yl)phenyl)acetamide (13 mg, 28%) as a white solid: MP = 208–211 °C; $^1$H NMR (300 MHz, DMSO-$d_6$) δ 9.98 (s, 1H), 8.00 (d, *J* = 9.3 Hz, 2H), 7.62 (d, *J* = 9.3 Hz, 2H), 7.51 (d, *J* = 1.5 Hz, 1H), 7.38–7.35 (m, 1H), 7.11 (d, *J* = 8.1 Hz, 1H), 6.81 (s, 1H), 4.05–3.99 (m, 1H), 3.95–3.93 (m, 1H), 2.30 (s, 3H), 2.27–2.25 (m, 2H), 2.04 (s, 3H); ESI MS *m/z* 350 [$C_{20}H_{19}N_3O_3$ + H]$^+$; HPLC (Method A) >99% (AUC), $t_R$ = 10.96 min; Chiral HPLC (Chiralpak AD, Method A) 97.6% (AUC), $t_R$ = 17.79 min.

#### 1.7. Preparation of (S)-3a-Hydroxy-6-methyl-1-(4-morpholinophenyl)-1,2,3,3a-tetrahydro-4H-pyrrolo[2,3-b]quinolin-4-one (Skeletostatin 3)

**Synthetic Scheme**

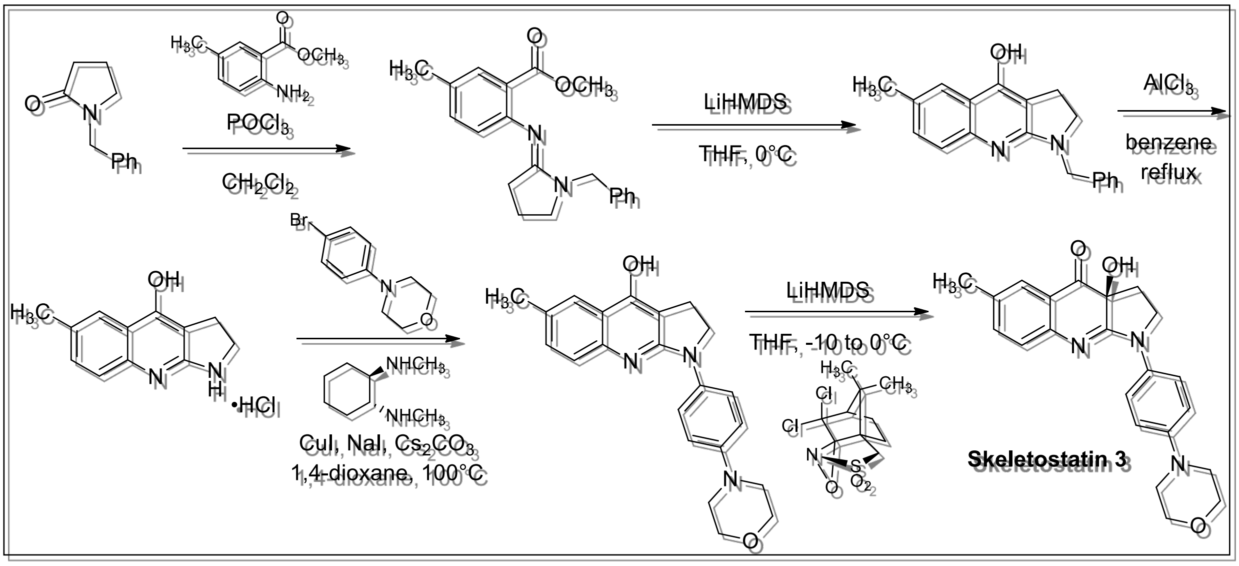

**Preparation of Methyl 2-((1-benzylpyrrolidin-3-ylidene)amino)-5-methylbenzoate**

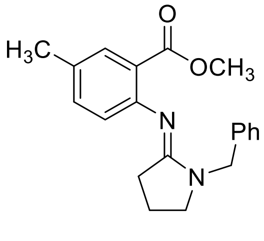

A solution of 1-benzylpyrrolidin-2-one (6.30 g, 35.9 mmol) in methylene chloride (200 mL) was treated with phosphorous oxychloride (4.50 mL, 49.2 mmol) and stirred under a nitrogen atmosphere at ambient temperature for 4 h. The mixture was treated with a solution of methyl 2-amino-5-methylbenzoate (5.30 g, 32.1 mmol) in methylene chloride (25 mL) and heated at reflux for 16 h. After this time, the reaction mixture was allowed to cool to ambient temperature and concentrated under reduced pressure. The residue was diluted with ethyl acetate and washed with saturated aqueous sodium bicarbonate. The organic layer was extracted with 1 N aqueous hydrochloric acid, and the aqueous layer was basified with 2 N aqueous sodium hydroxide. The mixture was extracted with ethyl acetate. The combined organics were washed with water and brine, dried over sodium sulfate, filtered, and concentrated under reduced pressure to provide methyl 2-((1-benzylpyrrolidin-3-ylidene)amino)-5-methylbenzoate (8.40 g, 81%): $^1$H NMR (300 MHz, $CDCl_3$) δ 7.59 (d, *J* = 1.8 Hz, 1H), 7.41–7.27 (m, 5H), 7.17 (dd, *J* = 7.8, 1.5 Hz, 1H), 6.75 (d, *J* = 8.1 Hz, 1H), 4.64 (s, 2H), 3.80 (s, 3H), 3.28 (t, *J* = 6.9 Hz, 2H), 2.34–2.29 (m, 5H), 1.95–1.85 (m, 2H); ESI MS *m/z* 323 [$C_{20}H_{22}N_2O_2$ + H]$^+$.

**Preparation of 1-Benzyl-6-methyl-2,3-dihydro-1*H*-pyrrolo[2,3-*b*]quinolin-4-ol**

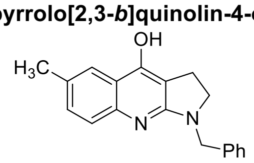

A solution of methyl 2-((1-benzylpyrrolidin-3-ylidene)amino)-5-methylbenzoate (8.40 g, 26.1 mmol) in tetrahydrofuran (200 mL) under a nitrogen atmosphere at 0 °C was treated dropwise with a 1.0 M solution of lithium bis(trimethylsilyl)amide in tetrahydrofuran (55 mL, 55 mmol) and stirred for 2 h. After this time, a saturated solution of ammonium chloride (100 mL) and water (100 mL) were added, and the mixture was stirred for 30 min. After this time, a precipitate formed. The precipitate was collected by filtration, washed with water and diethyl ether and dried under high vacuum to provide 1-benzyl-6-methyl-2,3-dihydro-1*H*-pyrrolo[2,3-*b*]quinolin-4-ol (3.18 g, 42%): ESI MS *m/z* 291 [$C_{19}H_{18}N_2O$ + H]$^+$.

**Preparation of 6-Methyl-2,3-dihydro-1*H*-pyrrolo[2,3-*b*]quinolin-4-ol Hydrochloride**

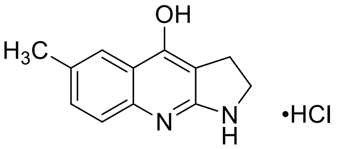

A mixture of 1-benzyl-6-methyl-2,3-dihydro-1*H*-pyrrolo[2,3-*b*]quinolin-4-ol (6.30 g, 21.7 mmol) and aluminum trichloride (12.8 g, 96.0 mmol) in benzene (140 mL) was stirred at reflux for 3 h. After this time, the mixture was cooled to room temperature and poured into stirred ice/water (250 mL). The mixture was stirred for 15 min and then the precipitate was collected by filtration, washed with diethyl ether and dried under high vacuum to provide 6-methyl-2,3-dihydro-1*H*-pyrrolo[2,3-*b*]quinolin-4-ol hydrochloride (5.1 g, quantitative): $^1$H NMR (500 MHz, DMSO-$d_6$) δ 13.51 (br s, 1H), 12.23 (br s, 1H), 8.75 (s, 1H), 7.80 (s, 1H), 7.54 (d, *J* = 8.5 Hz, 1H), 7.48 (dd, *J* = 8.0, 1.5 Hz, 1H), 3.81 (t, *J* = 8.0 Hz, 2H), 3.16 (t, *J* = 8.0 Hz, 2H), 2.41 (s, 3H); ESI MS *m/z* 201 [$C_{12}H_{12}N_2O$ + H]$^+$.

**Preparation of 6-Methyl-1-(4-morpholinophenyl)-2,3-dihydro-1*H*-pyrrolo[2,3-*b*]quinolin-4-ol**

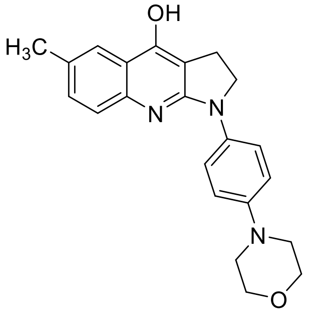

A solution 4-(4-bromophenyl)morpholine (2.01 g, 8.30 mmol) in 1,4-dioxane (20 mL) was treated with copper iodide (158 mg, 0.830 mmol), cesium carbonate (8.10 g, 24.9 mmol), sodium iodide (3.73 g, 24.9 mmol) and *N*,*N*'-dimethyl-(1*R*,2*R*)-1,2-cyclohexanediamine (235 mg, 1.66 mmol) and heated at 100°C for 16 h. After this time, the reaction was treated with copper iodide (80 mg, 0.42 mmol) and *N*,*N*'-dimethyl-(1*R*,2*R*)-1,2-cyclohexanediamine (100 mg, 0.706 mmol) and heated at 100°C for 64 h. After this time, the reaction mixture was cooled to ambient temperature. The mixture was diluted with ethyl acetate, washed sequentially with water, saturated aqueous ammonium chloride, water, and brine. The organic layer was dried over sodium sulfate, filtered, and concentrated under reduced pressure to provide a 7:3 mixture of 4-(4-iodophenyl)morpholine: 4-(4-bromophenyl)morpholine (1.89 g). A solution of 6-methyl-2,3-dihydro-1*H*-pyrrolo[2,3-*b*]quinolin-4-ol hydrochloride (280 mg, 1.18 mmol) in dimethyl sulfoxide (8 mL) was treated with a portion of the 7:3 mixture of 4-(4-iodophenyl)morpholine: 4-(4-bromophenyl)morpholine (550 mg), copper iodide (60 mg, 0.32 mmol), cesium carbonate (1.5 g, 4.60 mmol), and *N,N'*-dimethylethane-1,2-diamine (55 mg, 0.62 mmol) and heated at 160 °C in a sealed vial for 16 h. After this time, the reaction mixture was filtered through diatomaceous earth. The filtrate was concentrated under reduced pressure. The crude residue was purified by column chromatography (silica gel, 0–50% (methylene chloride/methanol/ammonium hydroxide 80/18/2)/methylene chloride) to provide 6-methyl-1-(4-morpholinophenyl)-2,3-dihydro-1*H*-pyrrolo[2,3-*b*]quinolin-4-ol (74 mg, 17%): $^1$H NMR (500 MHz, DMSO-$d_6$) δ 10.39 (br s, 1H), 7.81 (br s, 2H), 7.74 (s, 1H), 7.43 (d, *J* = 8.5 Hz, 1H), 7.27 (dd, *J* = 8.5, 2.0 Hz, 1H), 7.00 (d, *J* = 9.0 Hz, 2H), 4.01 (t, *J* = 8.0 Hz, 2H), 3.75 (t, *J* = 5.0 Hz, 4H), 3.11–3.07 (m, 6H), 2.39 (s, 3H); ESI MS *m/z* 362 [$C_{22}H_{23}N_3O_2$ + H]$^+$.

**Preparation of (*S*)-3a-Hydroxy-6-methyl-1-(4-morpholinophenyl)-1,2,3,3a-tetrahydro-4*H*-pyrrolo[2,3-*b*]quinolin-4-one**

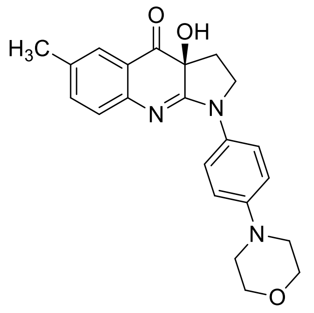

A solution of 6-methyl-1-(4-morpholinophenyl)-2,3-dihydro-1*H*-pyrrolo[2,3-*b*]quinolin-4-ol (70 mg, 0.19 mmol) in tetrahydrofuran (4 mL) at –10 °C was treated dropwise with a 1.0 M solution of lithium bis(trimethylsilyl)amide in tetrahydrofuran (0.23 mL, 0.23 mmol), followed by (-)-(8,8-dichlorocamphorylsulfonyl)oxaziridine (124 mg, 0.42 mmol) and stirred for 2 h. After this time, the mixture was treated with saturated aqueous ammonium iodide and stirred for 15 min. Saturated aqueous sodium thiosulfate was added, and the mixture was stirred for 15 min and then extracted with ethyl acetate. The organics were washed with water and extracted with 0.8 N aqueous hydrochloric acid, and the aqueous extract was basified with sodium hydroxide. The mixture was extracted with ethyl acetate, washed with water and brine, dried over sodium sulfate, filtered, and concentrated under reduced pressure. The crude residue was crystallized from hot acetonitrile to provide (*S*)-3a-hydroxy-6-methyl-1-(4-morpholinophenyl)-1,2,3,3a-tetrahydro-4*H*-pyrrolo[2,3-*b*]quinolin-4-one (27 mg, 38%) as a pale yellow-brown solid: mp = 226–227 °C; $^{1}$H NMR (500 MHz, DMSO-$d_6$) δ 7.91–7.88 (m, 2H), 7.50 (d, *J* = 2.0 Hz, 1H), 7.34–7.32 (m, 1H), 7.04 (d, *J* = 8.0 Hz, 1H), 7.02–6.98 (m, 2H), 6.76 (s, 1H), 4.05–4.00 (m, 1H), 3.92–3.88 (m, 1H), 3.75 (t, *J* = 5.0 Hz, 4H), 3.11 (t, *J* = 5.0 Hz, 4H), 2.29 (s, 3H), 2.25–2.22 (m, 2H); ESI MS *m/z* 378 $[C_{22}H_{23}N_3O_3 + H]^+$; HPLC (Method B) >99% (AUC), $t_R$ = 7.70 min; Chiral HPLC (Chiralpak AD, Method A) >99% (AUC), $t_R$ = 28.09 min.

### 2. Kinetic aqueous solubility.

#### 2.1. Materials

Verapamil hydrochloride and tamoxifen were obtained from Sigma Aldrich. All solvents were obtained from commercial sources and used without further purification. Hydrophilic PVDF 96-well filter plates (0.45 µm), 2 mL 96-well assay block, and 96-well UV plates were purchased from Fisher Scientific.

#### 2.2. Methods

<u>Kinetic aqueous solubility</u>

Test compound was prepared as a 10 mM stock solution in DMSO. Aqueous suspensions of test compound at 100 µM were prepared in PBS (0.01M) pH 7.4. The suspensions were agitated at 150 rpm for 1 hour at room temperature. The suspensions were then transferred to a 0.45 µm hydrophilic PVDF 96-well filter plate mounted on a fresh 2 mL 96-well plate and were filtered by centrifugation at 3,000 rpm for 1 min. 150 µL of filtrates were then transferred to 96-well UV plate for absorbance measurement.

<u>UV absorbance determination (for solubility assays)</u>

In order to determine the optimal wavelength for detection ($\lambda_{max}$), the absorbance spectrum of the DMSO stock solution for each compound was recorded over a broad range of wavelengths (200–700 nm) using a UV/VIS plate reader (Molecular Devices SpectraMax i3). To prepare calibration curves, a 1:1 serial dilution was performed on each compound starting at 80 µM to generate calibration solutions with concentrations ranging from 1.25 µM to 100 µM. The UV absorbance of calibrants was measured at $\lambda_{max}$ for each compound. Concentration of each compound in the assay solution (kinetic solubility) was calculated using a corresponding calibration curve. All measurements were done in duplicate.

### 3. Photostability.

#### 3.1. Materials

All solvents were obtained from commercial sources and used without further purification. Clear glass vials (2.0 mL with PTFE/silicone split septa) were purchased from Chemglass Life Sciences (CV-5326-1232).

#### 3.2. Methods

<u>Photostability</u>

Test compound was prepared as a 0.5 mg/mL stock solution in 1:1 acetonitrile/water. The stock solution (1.0 mL) was diluted with acetonitrile (1.0 mL) and water (2.0 mL) and transferred equally to four clear glass vials. One vial (t = 0 h) was stored in the dark at 4 ºC to await HPLC analysis. The remaining three vials were placed in a mirrored box with an LED light source (Nicrew 11W, 640 lumen aquarium light with white and blue LEDs, part number 6292452011) at ambient temperature. One of these vials was covered in aluminum foil to block out the light. After 4 hours, one of the clear vials was removed from the light box. After 24 hours, the remaining two vials (one clear and one foil-covered) were removed. All of the samples were analyzed by HPLC (254 nm) to determine the percentage of parent compound remaining.

### 4. Absorbance measurements

UV-VIS absorbance spectra were acquired on a NanoDrop 2000 spectrophotometer (Thermo Fisher Scientific) in the wavelength range of 190–840 nm at room temperature. Compounds were dissolved in DMSO at a concentration of 0.5 mM. The background (spectrum of pure DMSO) was subtracted from all spectra.

#### 5. Fluorescence spectroscopy

Fluorescence 3D scans were performed on a Cary Eclipse Fluorescence Spectrophotometer (Agilent Technologies). The excitation wavelength was changed from 250 nm to 550 nm in 10 nm increments while the emission wavelength was changed from 270 nm to 700 nm in 10 nm increments. The optical bandwidth was 10 nm on both the excitation and emission sides. Compounds were dissolved in dimethyl sulfoxide (DMSO) and diluted in Phosphate Buffered Saline (PBS) such that the final compound concentration was 5 µM for all blebb derivatives. In all samples, the final DMSO concentration was 1%. The background fluorescence of PBS containing 1% DMSO was subtracted from all spectra. Fluorescein at a final concentration of 5 nM was used as reference. All spectra were normalized to the highest peak detected in the spectrum of fluorescein (at $\lambda_{Ex}$ = 490 nm and $\lambda_{Em}$ = 510 nm).

#### 6. Purification and preparation of protein samples.

F-actin was prepared from rabbit muscle acetone powder (cat. 41995-2, Pel Freez Biologicals) as described by Pardee and Spudich[4]. Full length rabbit skeletal muscle myosin II (cat. MY02), full length bovine cardiac muscle myosin II (cat. MY03), and the S1 fragment of chicken gizzard smooth muscle myosin II (cat. CS-MYS05) were purchased from Cytoskeleton, Inc. Lyophilized protein samples were reconstituted following the manufacturer`s instructions. Full length NMIIA was isolated from human platelets and thiophosphorylated based on the modified protocol of Daniel and Sellers[5]. Briefly, freshly drawn human platelets from anonymous donors were obtained from the Department for Transfusion Medicine at NIH and homogenized in a Teflon-glass tissue grinder in 50 ml of extraction buffer containing 500 mM NaCl, 5 mM ethylenediaminetetraacetic acid (EDTA), 10 mM ethylene glycol-bis(β-aminoethyl ether)-N,N,N′,N′-tetraacetic acid (EGTA), 0.5 % TritonX-100, 5 mM adenosine triphosphate (ATP), 2 mM dithiothreitol (DTT), 0.1 mM phenylmethylsulfonyl fluoride (PMSF), 0.01 mg/ml leupeptin, 10 µM Cytochalasin D, 50 mM 3-(N-morpholino)propanesulfonic acid (MOPS, pH=7.0) and 1 SIGMAFAST Prot. Inh. Tablet (cat. S8820, Sigma). The homogenate was centrifuged for 30 min at 175,000 $g_{max}$. The supernatant was made 30 mM in $MgCl_2$ by adding 500 mM stock solution. An additional 5 mM ATP was added, while maintaining the pH between 6.8 and 7.0 with the addition of 1 M tris(hydroxymethyl)aminomethane (TRIS, base form). Saturated ammonium sulfate solution was slowly added with stirring to a final saturation of 20%. The solution was centrifuged for 10 min at 12,000 $g_{max}$. Subsequently, two other steps of ammonium sulfate precipitation were performed reaching final saturations of 40% and 60%. The final pellet was dissolved in “Buffer A” containing 20 mM NaCl, 10 mM $MgCl_2$, 0.1 mM EGTA, 1 mM DTT, 0.1 mM PMSF, 5 µg/ml leupeptin, and 10 mM MOPS (pH=7.0). The solution was dialyzed overnight against the same buffer to allow myosin to form filaments. Myosin filaments were sedimented by centrifugation at 40,000 $g_{max}$ for 20 min and the pellet was resuspended in a small amount of Buffer A. The actin contaminated crude myosin fraction was made 500 mM in NaCl to dissociate myosin filaments. The sample was incubated in the presence of 40 µM Cytochalasin D on ice overnight. (Cytochalasin D blocks actin polymerization, therefore, long incubations lead to decreased levels of actin filaments in the solution.) Subsequently, a second ammonium sulfate fractionation was performed without the addition of ATP (to allow myosin to bind actin). The NMIIA-Actin complex was precipitated below 45% $(NH_4)_2SO_4$. Contaminants precipitating between 45% and 60% were discarded. The actin contaminated crude myosin fraction was dissolved in “Buffer B” containing 500 mM NaCl, 0.1 mM EGTA, 1 mM DTT, 10 mM MOPS (pH=7.0). The sample was incubated in the presence of 10 µM Cytochalasin D on ice overnight. An additional 10 mM $MgCl_2$ and 5 mM ATP was added to the sample and third ammonium sulfate fractionation was performed, resulting in the efficient separation of actin and NMIIA. Actin precipitated at a saturation of 45%, while myosin precipitated between 45 and 60%. The pellet containing NMIIA was dissolved in Buffer B and dialyzed against the same buffer overnight. The regulatory light chain of NMIIa was thiophosphorylated by using rabbit smooth muscle myosin light chain kinase (MLCK) at a 1:300 kinase:substrate molar ratio in a buffer containing 1 mM adenosine 5′-O-(3-thiotriphosphate), 150 mM NaCl, 2.5 mM $MgCl_2$, 2.5 mM $CaCl_2$, 10 nM Calmodulin (cat. C-1001-1, rPeptide LLC), 10 mM MOPS (pH 7.2) at 25 °C for 3 hours. MLCK was a kind gift of Dr. James R. Sellers. The reaction was stopped by the addition of 5 mM EGTA. The NaCl concentration was increased to 500 mM. Phosphorylated NMIIa was isolated from the reaction mixture by ammonium-sulfate precipitation. The protein precipitated between 40 and 60% $(NH_4)_2SO_4$. The pellet was dissolved in Buffer B and dialyzed against the same buffer overnight. Myosin light chain thiophosphorylation was confirmed by urea polyacrylamide gel electrophoresis as described by Daniel and Sellers[5].

#### 7. NADH-coupled ATPase assays.

ATPase assays were performed using a nicotinamide adenine dinucleotide (NADH)-coupled ATPase assay as previously described[6]. In this assay, the hydrolysis of ATP is coupled to the oxidation of NADH by enzymatic reactions. Therefore, the ATP consumption rate can be measured by following the decrease in NADH concentration based on the intrinsic fluorescence of NADH. Experiments were performed at 25 °C in 384-well plates in a total reaction volume of 20 µl. The concentration of full length SkMII, CMII, and NMIIA was 20 nM, 300 nM, and 400 nM, respectively, in all experiments. Concentration of the S1 fragment of SmMII was 400 and 200 nM in the case of blebb and skeletostatin 1, respectively. The concentration of actin was 10 µM in all experiments. To determine inhibitory constants, ATP consumption rates were plotted against the inhibitor concentration and the dose-response data were fitted to a quadratic equation corresponding to a simple one-to-one binding equilibrium model[6].

### 8. Cytokinesis assay.

The cytokinesis assay was performed as previously described[7]. Briefly, COS7 cells were plated at 2000 cells per well in 96-well plates and grown at 37 °C for 24 hours. NMIIB is expressed almost exclusively among other NMII heavy chains in COS7 cells[8]. Therefore, cytokinesis (which is an NMII-dependent process) depends almost entirely on the activity of NMIIB in these cells. Compounds were dissolved in DMSO and dilution series were prepared as required. Right before treating cells, compound solutions were diluted fifty-fold into cell growth medium. Equal volumes of the diluted compound solutions and cell cultures were combined, resulting in another two-fold dilution and a final DMSO concentration of 1%. Treated cell cultures were incubated for 24 hours, stained with a combination of fluorescein-diacetate, Hoechst 33342, and propidium iodide and imaged. (These dyes stain living cell bodies, nuclei and nuclei of dead cells, respectively.) Images were analyzed by counting cells and nuclei and determining the nuclei to cell ratio (NCR) for each well. In the presence of cytokinesis inhibitors, this ratio increases over time, as the ratio of multinucleated cells increases in the population. (Nuclei can divide, but the separation of daughter cells fails.) Compound potencies ($EC_{50}$) were determined by fitting the dose-response data to the Hill-equation. The ratio of dead nuclei to total nuclei provided a simultaneous measure of cytotoxicity[7].

### 9. *In vivo* studies in mice

#### 9.1. Animals.

Adult, 8-10 week old male and female C57BL/6 mice (25-30 g, Jackson Laboratory) were housed under a 12:12 light/dark cycle, with food and water ad libitum. All procedures were performed in accordance with the Scripps Research Institutional Animal Care and Use Committee. Mice were handled for 3-5 days before behavioral testing.

#### 9.2. Open field.

To examine the effect of Skeletostatin 1 on general locomotion, mice were injected with either vehicle, 2 mg/kg, 5 mg/kg, or 10 mg/kg (IP) Skeletostatin 1 and allowed to freely explore an open field box for 5 or 60 min. Skeletostatin 1 was formulated for IP administration by first preparing a 30% Hydroxypropyl-β-Cyclodextrin (“HPβCD”, cat. 50592055, Fisher Scientific) solution (30 g HPBCD dissolved in 70 ml acetate buffer (100 mM, pH=5.4). The HPβCD solution was filtered using a 0.2 µm nylon syringe filter (cat. 09719C, Fisher Scientific). Subsequently, Skeletostatin 1 stock solutions at a concentration of 4.5 mg/ml, 11.25 mg/ml, and 22.5 mg/ml were prepared in pure DMSO (cat. D2650, Sigma). Right before administration, 20 µl Skeletostatin 1 stock solution was added to 280 µl HPβCD solution in a 1.5 ml microcentrifuge tube, yielding a final DMSO concentration of 6.67% (v/v) and final dose concentrations of 0.3 mg/ml, 0.75 mg/ml, and 1.5 mg/ml, respectively. Solutions were mixed vigorously by shaking, then administered to mice (IP) at a dose volume of 6.67 ml/kg yielding dose levels of 2 mg/kg, 5 mg/kg, and 10 mg/kg, respectively. Immediately after injection, mice were placed in the open field apparatus. Open field arenas consisted of custom-made clear acrylic boxes (43 x 43 x 32h cm) with opaque white acrylic siding surrounding each box (45 x 45 x 21.5h cm). Activity was monitored with CCTV cameras (Pansonic WV-BP334) feeding into a computer equipped with Ethovision XT (Noluds Information Technology) for data acquisition and analysis. Behavior was recorded and analyzed using Ethovison software. Immediately following the end of the session, mice were euthanized, and tissue and plasma were collected to determine Skeletostatin 1 concentrations (see Pharmacokinetics section).

#### 9.3. Pharmacokinetics.

Mice were anesthetized with isoflurane (cat. IsoSol, Patterson Veterinary Supply Inc) and euthanized at 5 or 60 min post-injection. Blood samples were collected immediately into lithium heparin coated tubes (cat. 07 6101, Ram Scientific), and stored on wet ice. Blood samples were later centrifuged for 3 min at 5,000 RMP to separate plasma from red blood cells. Plasma was collected into a fresh tube and stored at -80 °C. In addition to blood samples, skeletal muscle was collected from each mouse from their left hind thigh muscle. Each sample was flash frozen with 2-methylbutane and stored at -80 °C. Compound levels were quantified in muscle and plasma by mass spectrometry using an ABSciex 5500 mass spectrometer using multiple reaction monitoring. Muscle samples were homogenized in saline and then immediately treated with 2-times (v:v) acetonitrile to extract the compound and precipitate cellular protein. Plasma samples were directly treated with acetonitrile. Samples were filtered through a 0.2 µm filter plate prior to injection onto the LC-MS/MS. HPLC and MS/MS parameters:

LC (Shimadzu UFLC XR) conditions

| Compound | BPN-0027377 | I.S. (Carbamazepine) |
|---|---|---|
| Column | Thermo Betasil C18 5µ, 50x2.1mm | |
| Mobile phase | A: Water with 0.1% Formic Acid<br>B: Acetonitrile with 0.1% Formic Acid | |
| Flow rate (ml/min) | 0.35 | |
| Temperature (°C) | 35 | |
| Injection volume(µl) | 10 | |
| RT(min) | 2.3 | 2.5 |

Gradient elution conditions:

| Time (min) | Mobile phase A (%) | Mobile phase B (%) |
|---|---|---|

| | | |
|---|---|---|
| 0.2 | 90 | 10 |
| 0.5 | 90 | 10 |
| 2.0 | 5 | 95 |
| 3.0 | 5 | 95 |
| 4.0 | 90 | 10 |
| 5.9 | 90 | 10 |

MS (API5500) conditions

| Compound | BPN-0027377 | I.S. (Carbamazepine) |
|---|---|---|
| MRM(+) | 349.1/162 | 237.2/194.1 |
| Collision Gas | 7 | |
| Curtain GAS | 36 | |
| Ion Source Gas1 | 55 | |
| Ion Source Gas2 | 50 | |
| Ion Spray Voltage | 5500 | |
| Temperature (°C) | 550 | |
| Collision Energy | 21 | 26 |
| Declustering Potential | 31 | 136 |
| Entrance Potential | 10 | |
| Collision Cell Exit Potential | 14 | |

**9.4. Rotarod.**

Rotarod was used to further assess Skeletostatin 1 effects on motor function at the higher doses (5 and 10 mg/kg). Dosing was performed as described above (see "Open field"). Control mice were injected with vehicle only (6.67% (v/v) DMSO in HPβCD solution). The performance of mice was tested using a five-lane accelerating rotarod (Med Associates) in two consecutive trial sessions separated by 40 min intervals. The latency to fall off the rod (the time spent on the rod by each mouse) was recorded as the rod accelerated from 4 to 40 rpm over 5 min.

**9.5. Data analysis**

All behavioral experiments were analyzed using a one-way Analysis of Variance (ANOVA) or independent *t*-test, where applicable. All *post hoc* tests were conducted, when appropriate, using Tukey's Honest Significance Test (HSD) test.
